# Phase Separation Tunes the Stability and Dynamics of G-quadruplex and i-Motif DNA in the Nuclei of Living Cells

**DOI:** 10.1101/2025.06.30.661982

**Authors:** Brahmmi Patel, Cailin Hoang, Hyejin Yoo, Caitlin Davis

## Abstract

G-quadruplexes and i-motifs are non-canonical DNA structures that are important not only as targets of gene regulation in cancer therapy, but also as nanotools with biomedical applications. However, their apparent in vitro stabilities and dynamics cast doubt upon their physiological relevance and utility *in vivo.* Here we investigate folding of the thrombin-binding aptamer G-quadruplex and the TAA(C_5_TAA)_4_ i-motif. We use fast relaxation imaging, which couples fluorescence microscopy with laser-induced temperature jumps, to quantify the stability and dynamics of FRET-labelled constructs in vitro and in the nuclei of living cells. Our work suggests that the stability and kinetics of i-motif and G-quadruplex DNA are carefully tuned by phase separation to fit the seconds-to-minutes timescales of key regulatory processes inside cells. Taken together, our findings underscore the importance of studying the folding of DNA structures under near-physiological conditions to their mechanistic understanding as well as for the development of DNA-targeted drugs.

**GRAPHICAL ABSTRACT:** 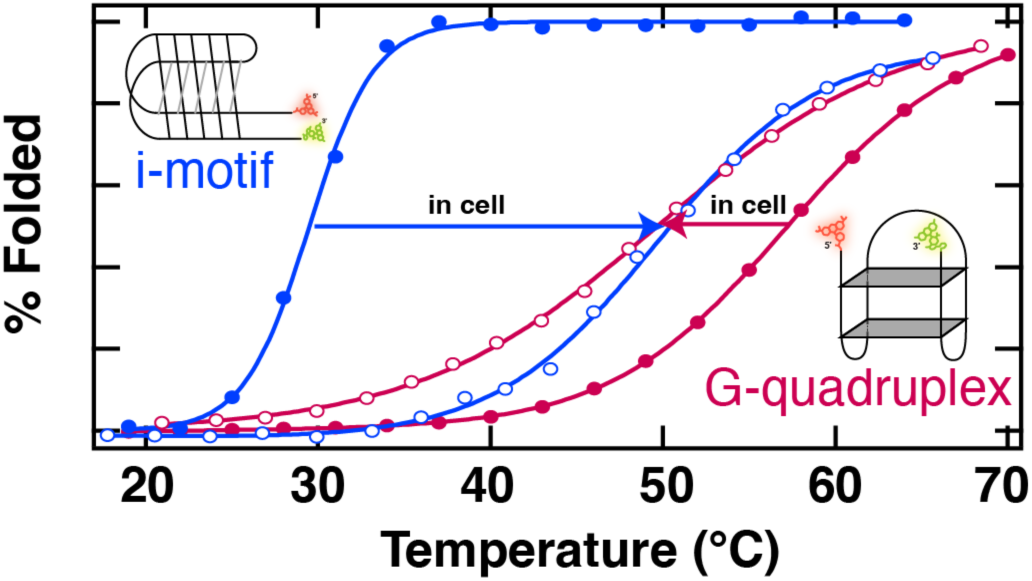

## INTRODUCTION

The stability and dynamics of G-quadruplex and i-motif DNA are crucial for their use in biomedical, pharmacological, and nanotechnological applications. G-quadruplexes and i-motifs are non-canonical DNA structures found in complementary guanine-rich and cytosine-rich stretches of the genome, respectively, with a high prevalence in promoter and telomeric regions of DNA, including many oncogenes.^1–3^ Moreover, it is the presence of these structures, rather than the underlying DNA sequence, that drives changes in gene expression and the chromatin landscape.^4–7^ However, the apparent stabilities and dynamics of these non-canonical DNA structures under physiological conditions in vitro have led to skepticism towards their biological relevance. G-quadruplexes tend to have thermal stabilities far above physiological (37 °C) that are further stabilized by environmental conditions anticipated in cells, such as high ionic strength and macromolecular crowding.^8,9^ Conversely, most i-motifs do not fold at neutral pH in dilute aqueous solution, and the effects of macromolecular crowding are modest.^10^ Due to these observations, their existence in vivo remained speculative until recently when the presence of folded G-quadruplexes and i-motifs inside living cells was confirmed by small molecule or antibody binding.^11–14^ However, the folding of G-quadruplexes and i-motifs in vitro has not been quantitatively compared to living cells. Understanding their stabilities and dynamics in the native cellular context is critical for the advancement of targeted therapies and drug delivery systems in living cells and has far-reaching implications in the biomedical use of these structures.

Here, we measure the stability and dynamics of a FRET-labelled G-quadruplex and i-motif DNA in the nuclei of living cells. To interpret our in-cell measurements we compare with analogous in vitro experiments in dilute buffer and cellular mimetics. We selected the thrombin-binding aptamer (TBA) as our G-quadruplex model since it is well-characterized in vitro with a near physiological thermal stability, 50 °C.^15^ The TBA G-quadruplex (Figure 1) is formed by two stacked coplanar guanine tetrads held together by Hoogsteen hydrogen bonding and coordinated by a central monovalent cation. TBA has potential for many biomedical applications due to its inhibition of thrombin protease, the key enzyme in coagulation.^16^ We selected the d[CCCTAA]_4_ human telomeric repeat as our model i-motif as it is formed in telomeric DNA in vivo and forms an intramolecular structure that is well characterized.^17^ To enable comparison of in vitro and in cells, the stability of the i-motif was raised by increasing the length of the C-tract, TAA(C_5_TAA)_4_.^2,18,19^ The TAA(C_5_TAA)_4_ i-motif (Figure 1B) is assembled from four parallel cytosine-rich strands held together by hemi-protonated intercalated cytosine (C:C+) base pairs. The human telomeric repeat i-motif is a promising target for anticancer drug design due to its roles in telomere function and telomerase activity regulation.

**Figure 1:**
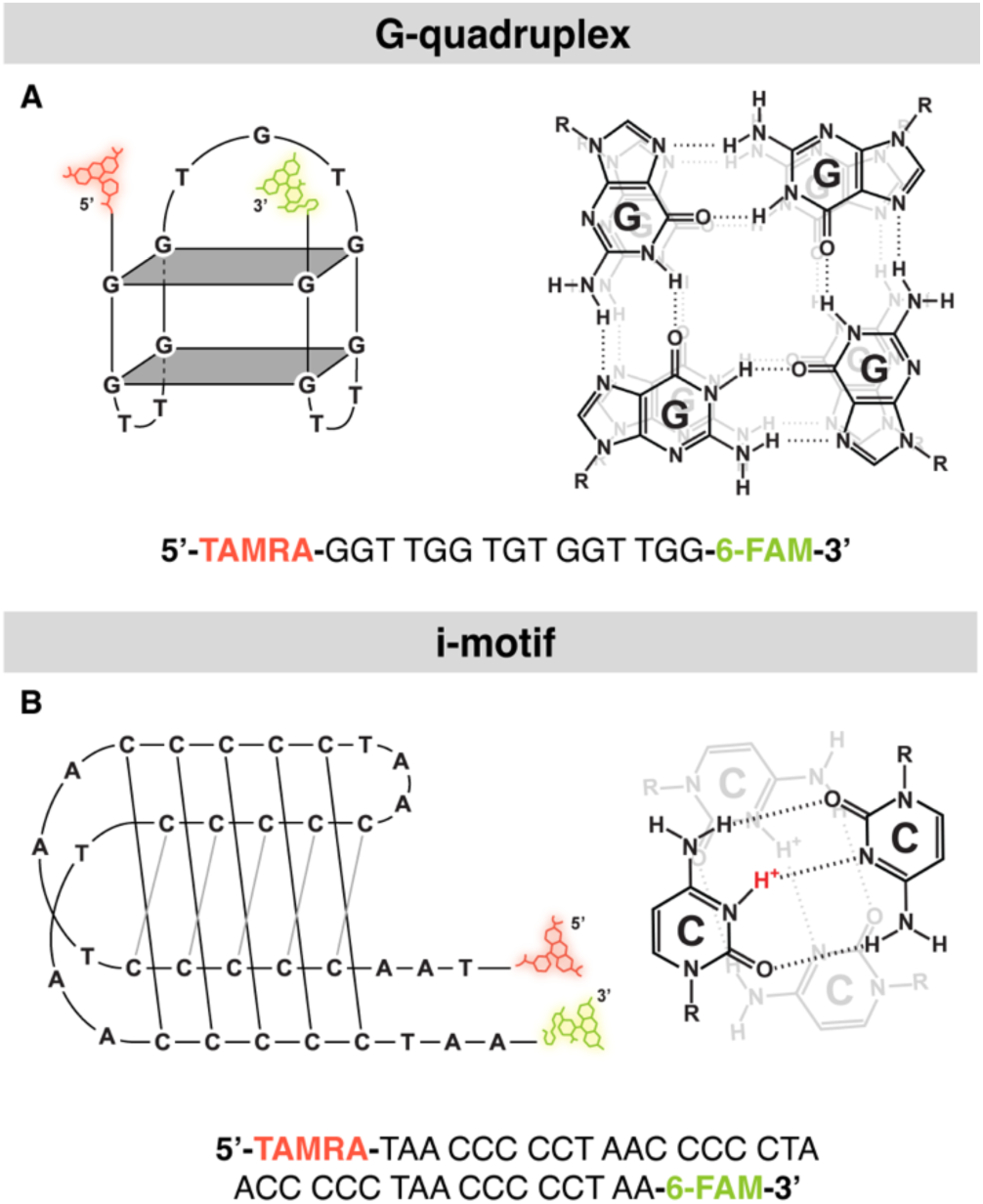
G-quadruplex and i-motif DNA models are used for in vitro and in cell comparison of stability and kinetics. Cartoon of the secondary structure (left), chemical structure (right), and oligonucleotide sequence (bottom) of the FRET-labelled (A) TBA G-quadruplex DNA and (B) TAA(C^5^TAA)4 i-motif DNA. Image generated in Adobe Illustrator.

To quantify thermodynamics and kinetics of the g-quadruplex and i-motif in vitro and in living cells, we used Fast Relaxation Imaging (FReI), which combines fluorescence microscopy of FRET labeled biomolecules with laser-induced temperature jumps. In-cell FReI has been previously used to study folding of proteins and RNA as well as to quantify binding of proteins and nucleic acids.^20–23^ Using our FRET-based approach, we quantify the in vitro and intranuclear stability and kinetics of both the FRET-labeled TBA G-quadruplex (G4) and FRET-labeled TAA(C_5_TAA)_4_ human telomeric repeat i-motif (iM). The intranuclear effects on the DNA are dramatic. While G4 is destabilized by ≈10 °C inside cells compared to in vitro, iM is stabilized by nearly 20 °C. These observations cannot be explained by macromolecular crowding or non-specific chemical interactions. Instead, we find that specific chemical interactions with proteins inside the cell nuclei are responsible for the opposing changes in stability of G4 and iM. The folding kinetics of G4 and iM are also oppositely affected: inside cells, G4 folding is slowed 36x while iM folding is sped 16x. We find that the kinetics of G4 and iM inside cells are primarily governed by phase separation triggered by interactions with positively charged species in the nuclear environment. No single in vitro condition could reproduce both the intranuclear stability and kinetics of the DNA. Taken together, our work emphasizes the importance of studying non-canonical DNA directly in cells, especially for biomedical applications where the effectiveness of therapies relies on targeting the correct ensemble of states.

## MATERIAL AND METHODS

### DNA Oligonucleotides

TBA G-quadruplex (5’ – GGT TGG TGT GGT TGG – 3’) and TAA(C_5_TAA)_4_ human telomeric repeat i-motif mutant (5’ – TAA CCC CCT AAC CCC CTA ACC CCC TAA CCC CCT AA – 3’) were purchased from Integrated DNA Technologies (IDT, Coralville, IA). To enable direct comparison between in vitro and in-cell measurements, the DNA were FRET-labelled (IDT) at the 5’ and 3’ ends with TAMRA (NHS ester) and 6-FAM (Fluorescein), respectively. For the colocalization experiments, the DNA sequences were labelled with Cy5.5 (NHS ester) at the 3’ end were purchased from IDT. Table S7 contains a complete list of oligos used in the study. Stock solutions of oligos were prepared at 100 µM in UltraPure DNase/RNase-Free Distilled Water (ThermoFisher Scientific, Waltham, MA) and stored at -20 °C.

### Buffer Preparation

Polyethylene glycol 10,000 (PEG10k, Sigma-Aldrich, St. Louis, MO) was dissolved at 100-400 mg/ml in filtered potassium phosphate buffer (10 mM potassium phosphate, 200 mM KCl, 1 mM K_2_EDTA, pH 7.0) and rocked overnight. To ensure that PEG was completely dissolved, the solution was heated to 55 °C for 20 minutes and cooled to room temperature prior to use.

Other buffers are commercially available. Nuclear Extraction Reagent (NER, ThermoFisher Scientific) was diluted to the desired concentration in potassium phosphate buffer. DPBS (Gibco, Grand Island, NY) was diluted from the manufacturer’s 10X stock to 1X in distilled water, pH adjusted to 7.0 and filtered prior to use. Salmon sperm DNA (ssDNA, ThermoFisher Scientific) was dialyzed overnight at room temperature in phosphate buffer (10 mM potassium phosphate, 200 mM KCl, 1 mM K_2_EDTA, pH 7.0). A mixture of 10 mg/ml BSA (Sigma), 37.5 %v/v NER, 2.5 mg/ml ssDNA, and 150 mg/ml PEG10k in 10 mM phosphate buffer pH 7 was prepared from buffer stocks.

Prior to in vitro analysis, the oligo stock was diluted to the final working concentration in the desired experimental buffer and annealed in a Nexus Gradient Thermal Cycler (Eppendorf, Hamburg, Germany); following established protocols,^24^ each sample was heated to 90 °C for 5 minutes before slow cooling to 4 °C at a rate of 0.5 °C/min. To avoid refolding ssDNA, oligo stocks were annealed in the thermocycler prior to dilution in buffers containing ssDNA.

### Nuclear Cell Lysate Preparation

Cultured human bone osteosarcoma epithelial cells, U-2 OS (ATCC HTB-96, Manassas, VA), were grown to ≈80% confluency in a T175 cell culture flask (ThermoFisher, Scientific). DMEM (Corning, Corning, NY) + 10% fetal bovine serum (FBS, ThermoFisher Scientific) + 1% penicillin-streptomycin (P/S, Corning) media was aspirated, and cells were washed with 10 ml of 1X PBS buffer (ThermoFisher Scientific). PBS was aspirated and 1 ml of pre-warmed trypsin-EDTA (Corning) was added. The flask was placed in a 37 °C incubator for 5 minutes and then resuspended in 10 ml of 4 °C PBS buffer to inactivate the trypsin. The cell suspension was spun down at 500 xg for 3 minutes. The pellet was rinsed with 10 ml of PBS buffer before being spun down again at 500 xg for 3 minutes. Nuclear cell lysate was prepared using the NE-PER^TM^ Nuclear and Cytoplasmic Extraction Reagents Kit (ThermoFisher Scientific) and following the manufacturer’s protocol. The nuclear cell lysate was dialyzed overnight in phosphate buffer (10 mM potassium phosphate, 200 mM KCl, 1 mM K_2_EDTA, pH 7.0) at room temperature in a Slide-A-Lyzer^TM^ 3.5K MWCO MINI Dialysis Device (ThermoFisher Scientific). The concentration of the nuclear lysate was quantified using a Pierce^TM^ Bradford Protein Assay Kit (ThermoFisher Scientific) and used immediately. To avoid denaturing proteins in the lysate, oligo stocks were annealed in the thermocycler prior to dilution in nuclear cell lysate.

### Turbidity Assay

Turbidity was assessed in solutions with increasing concentrations of DNA and 0.1 mg/ml unlabeled poly-L-lysine (PLL). 5 μM of pre-annealed unlabeled DNA was diluted to the desired final concentration in a solution containing 0.1 mg/ml unlabeled PLL (25988-63-0, Electron Microscopy Sciences) or FITC-labeled PLL (P3069, Sigma) in phosphate buffer (10 mM potassium phosphate, 200 mM KCl, 1 mM K_2_EDTA pH 7.0) and incubated for 30 minutes at RT. After incubation, the absorbance at 400 nm of the unlabeled mixture was measured using a Nanodrop One spectrophotometer (Thermo Scientific). An average of three readouts was recorded. Fluorescent samples were used for complementary fluorescence imaging.

### Colocalization

Pre-annealed 5 μM G4-Cy5.5 or iM-Cy5.5 was incubated with 0.1 mg/ml FITC-PLL (P3069, Sigma) in phosphate buffer (10 mM potassium phosphate, 200 mM KCl, 1 mM K_2_EDTA pH 7.0) for 30 minutes at RT. To excite the FITC fluorophore, a white LED (X-Cite mini+, Excelitas, Waltham, MA) is passed through an MF475/35x bandpass filter (Thorlabs, Newton, NJ) and then reflected onto the sample by a DMLP505R dichroic mirror (Thorlabs). Fluorescence emission is passed through a long-pass filter (DMLP505, Thorlabs) and split into red and green light by an Optosplit III emission image splitter (Cairn Research, Kent, UK) containing a dichroic mirror (DMLP505R, Semrock). To excite the Cy5.5 fluorophore, the white LED (Excelitas) is passed through an MDF-Cy55 bandpass filter (Thorlabs) and then reflected onto the sample by an MDF-Cy55 dichroic mirror (Thorlabs). Fluorescence emission is passed through a long-pass filter (FELH0700, Thorlabs) and split into the two channels by the Optosplit containing a dichroic mirror (MDF-Cy55 dichroic, Thorlabs). Images are collected at a frame rate of 1 fps.

### Fluorescence Spectroscopy

Steady-state FRET measurements were performed on a FP-8500 Spectrofluorometer (JASCO, Easton, MD) equipped with an automatic 4-position Peltier cell changer with temperature controller (PCT-818, JASCO). Experiments were performed at 0.50 μM in a 10 mm pathlength microcuvette (16.160F-Q-10/Z15, Starna Cells, Atascadero, CA). 6-FAM was excited at 480 nm with a 5 nm slit width, and fluorescence was monitored from 475 to 700 nm in 1 nm intervals. Thermal denaturation was performed from 10 to 79 °C in 3 °C intervals with an equilibration time of 3 minutes prior to acquisition. Samples were covered with mineral oil to prevent evaporation.

### DNA Microinjection and Transfection

DNA was introduced into U-2 OS cells via microinjection or transfection. U-2 OS cells were cultured and grown to 80% confluency on coverslips (No. 1.5, MatTekCorp., Ashland, MA) in DMEM containing 1% P/S and 10% FBS. For microinjection, cells were rinsed with PBS and resuspended in Fluorobrite (Gibco) supplemented with 10% FBS (FemtoJet 4i, Eppendorf, Enfield, CT). The 100-200 μM stock of FRET-labeled oligos was annealed in a thermocycler before microinjection into cells. Alternatively, cells were transfected with a 1:3 µg:µl ratio of DNA plasmid to JetPRIME^®^ Transfection Reagent (Polyplus, Illkirch, FR) and incubated at

37 °C for 16 hours before imaging. Microinjected or transfected cells were rinsed with PBS, resuspended in Fluorobrite supplemented with 10% FBS, and mounted onto a microscope slide (1 mm, VWR, Radnor, PA) with a 120 μm spacer (GraceBioLabs, Bend, OR) for FReI imaging.

### FReI

The FReI setup has been described previously.^22^ Briefly, a 1950 nm infrared laser (AdValue Phototonics, Tucson, AZ) introduces a rapid step-function shaped temperature perturbation to the sample after which the sample is given 10-180 s to equilibrate. The temperature change is determined using the temperature-dependence of the quantum yield of mCherry.^25^

Fluorescence from the laser-heated field of view is imaged through an objective lens (EC Plan-Neofluar 63X/0.95, Zeiss, Oberkochen, Germany) onto a CMOS camera (Moment, Teledyne Photometrics, Tucson, AZ). To excite the 6-FAM donor fluorophore, a white LED (X-Cite mini+, Excelitas, Waltham, MA) is passed through an ET480/40X bandpass filter (Chroma, Bellows Fall, VT) and then reflected onto the sample by a FF509-FDi 01 dichroic mirror (Semrock, West Henrietta, NY). Fluorescence emission is passed through a long-pass filter (ET510lp, Chroma) and split into red and green light by an Optosplit III emission image splitter (Cairn Research, Kent, UK) containing a dichroic mirror (FF560-FDi 01, Semrock).

Images are collected at a frame rate of 5 fps for in-cell measurements and 1-40 fps for in vitro measurements.

### Analysis of Thermodynamic Data

All thermal denaturation curves were fit to a two-state equilibrium model^20^:

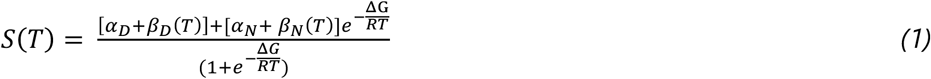

where *α_N_* and *α_D_* are the signals from the native (N) and denatured (D) states, respectively, and *β_N_* and *β_D_* are the slopes of the baselines from the native and denatured state, respectively. *R* is the ideal gas constant and *ΔG* is the free energy of unfolding (N to D) and can be approximated as a linear function of temperature: *ΔG* ≈ *ΔG*_1_(*T* – *T*_m_).^20^

### Analysis of Kinetic Data

The kinetics were plotted as the donor/acceptor ratio (D/A) versus time and fit to a double exponential function:

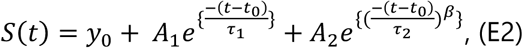

where the signal, S, as a function of time, *t*, is described by the sum of two exponential functions with pre-exponential factors, *A*_1_ and *A*_2_, and relaxation lifetimes, *1*_1_ and *1*_2._ The first exponential accounts for the instrument response, relaxation lifetimes of the fluorophores, and fast unresolved dynamics of the system. The second stretched exponential describes the folding/unfolding dynamics of the system following the temperature jump where *1*_2_ is the relaxation lifetime. The stretched exponential factor, *β*, allows for deviations from a simple two-state folding mechanism.^26^

## RESULTS

### G4 and iM are folded near physiological temperature and pH in crowded environments

Before investigating the stability of G4 and iM inside cells, we performed equilibrium circular dichroism measurements in vitro on labeled and unlabeled constructs to validate that FRET-labeling minimally perturbs their structure and thermal stability (SI Methods and Discussion, Figure S1). FRET has been used previously to study the in vitro folding of G-quadruplex and i-motif DNA.^27,28^ A donor (D) and acceptor (A) fluorophore are covalently attached to the 5’ and 3’ ends of the DNA, and FRET reports on the end-to-end distance of the DNA. A low donor/acceptor (D/A) ratio occurs when there is high FRET, where the fluorophores are close together and energy is transferred non-radiatively from the excited donor fluorophore to the acceptor fluorophore, such as in the folded state. A high D/A ratio indicates that the fluorophores are farther apart, such as in the unfolded state, reducing the efficiency of the energy transfer.

To predict the nuclear behaviors of G4 and iM, we investigated their folding in vitro near the physiological pH and macromolecular crowding concentration present in the cell (pH 7.0 and 200 mg/ml PEG10k).^29,30^ The fluorescence spectroscopy detected temperature-induced denaturation of G4 and iM is shown in Figure S2. The in vitro melting temperatures of G4 and iM are 58.9 ± 0.2 °C and 28.6 ± 0.7 °C, respectively (Figure 2). Ideal constructs for in cell folding studies will have stabilities at near-physiological temperatures that are not harmful to the cells. The in vitro stability of G4 is at the temperature limit of what has been previously measured in cells; infrared heating has been used to study folding as well as heat-shock response and gene induction up to 60 °C.^22,31^ Therefore, the in-cell stability of G4 was measured before making any destabilizing mutations to the sequence.

**Figure 2:**
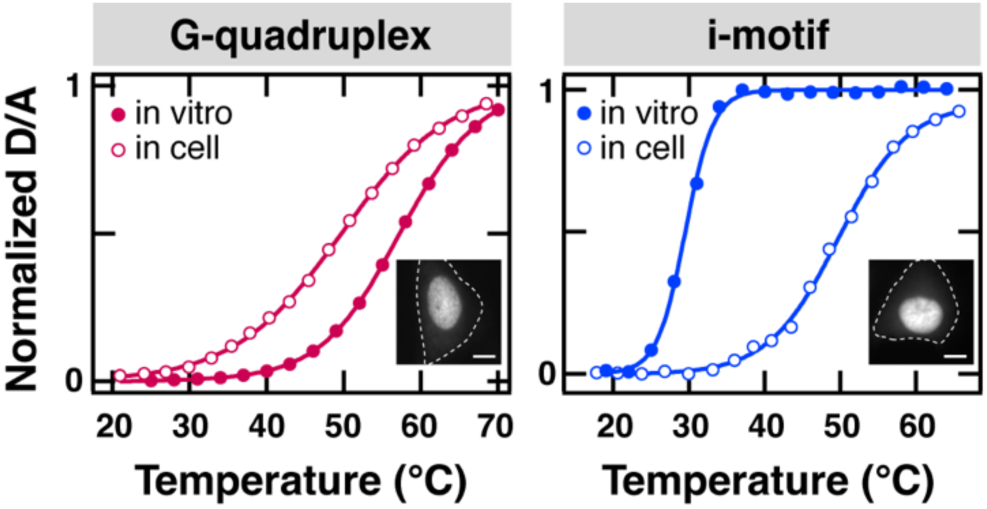
G4 is destabilized and iM is stabilized inside living cells. Representative equilibrium temperature denaturation of **(A)** G4 and **(B)** iM monitored by fluorescence spectroscopy in vitro and FReI microscopy in the nucleus of U-2 OS cells. In vitro experiments were conducted in 200 mg/ml PEG10k (10 mM potassium phosphate, 200 mM KCl, 1 mM K^2^EDTA, pH 7.0). The temperature dependent D/A ratios are fit (continuous line) to a two-state equilibrium model (Equation 1) and normalized to correct for pre- and post-transition baselines. Inset: Representative U-2 OS cells microinjected with FRET-labelled oligo that is localized to the nucleus. Cell outlined by white dashed line. Scale bar is 5 μm.

### G4 is destabilized whereas iM is stabilized by the nuclear environment

The stabilities of G4 and iM in the nucleus of living cells were measured by FReI. FReI combines fluorescence microscopy of FRET labeled biomolecules with small mid-IR laser-induced temperature jumps to thermally perturb the folding equilibrium. Thermal denaturation induced by the laser is monitored by changes in the donor and acceptor channels, and the stability is reported using the D/A ratio after the sample has time to fully equilibrate to the higher temperature. This approach is well established to study the folding of proteins and nucleic acids inside living cells with high spatiotemporal resolution.^21,22,32,33^ Figure 2 shows representative FReI melting curves for G4 and iM in the nucleus of living cells.

Inside the nucleus, G4 has an average melting temperature of 44.9 ± 0.9 °C (n=36) (Figure 2A, Table S1) compared to 58.9 ± 0.2 °C predicted by the in vitro FRET melts. The stability of G-quadruplexes is highly dependent on the identity of chemical species in solution.^34^ For example, G-quadruplexes are more stable in buffers containing potassium ions than buffers containing sodium ions.^9^ Indeed, at neutral pH, G4 is 25 °C less stable in DPBS, which is predominantly Na^+^, than in potassium phosphate buffer (SI Discussion, Figure S3A, Table S2).

Thus, the 14 °C decrease in stability is likely due to destabilizing chemical interactions with solutes, small molecules, and macromolecules in the nucleus of the cell.

Whereas G4 is destabilized inside cells, iM is significantly stabilized. iM has an average melting temperature of 46.5 ± 0.6 °C (n=60) inside the nucleus (Figure 2B, Table S3) compared to 28.6 ± 0.7 °C predicted by in vitro fluorescence melts. The 18 °C stabilization in the nucleus of cells could arise from macromolecular crowding, which is always stabilizing, or chemical interactions, which can be stabilizing or destabilizing. Like G4, iM is destabilized in sodium buffers when compared to phosphate buffers (Figure S3B, Table S4). Therefore, further experiments are necessary to determine the origin of differences in the stability of G4 and iM in the nucleus of cells.

### Nuclear measurements cannot be explained by macromolecular crowding alone

In cell measurements demonstrate that our near physiological buffer (pH 7.0 and 200 mg/ml PEG10k) is insufficient to replicate the nuclear environment. The total nuclear concentrations of macromolecules in eukaryotic cells range from 100 to 200 mg/ml.^29^ Therefore, to better understand the role that macromolecular crowding in the nucleus plays on folding of G4 and iM, their stabilities were measured in the presence of PEG10k between 0 and 300 mg/ml (Table 1, Figure S4). Macromolecular crowding theory predicts that the excluded volume induced by large macromolecules will result in entropic destabilization of extended or unfolded conformations, resulting in more energetically favorable compact, folded states.^35^ Indeed, the thermal stability of both oligos increased with increasing concentrations of PEG10k.

**Table 1:** Melting temperatures T_m_ extracted from temperature-dependent fluorescence spectroscopy measurements of G4 and iM in increasing concentrations of PEG10k (10 mM potassium phosphate, 200 mM KCl, 1 mM K_2_EDTA, pH 7.0). Error reflects the standard error of three replicates.

| [PEG 10k] mg/ml | G-quadruplex $T_m$ (°C) | i-motif $T_m$ (°C) |
| --- | --- | --- |
| 0 | $54.6 \pm 0.2$ | $27.5 \pm 0.3$ |
| 100 | $56.0 \pm 0.2$ | $28.5 \pm 0.7$ |
| 200 | $58.9 \pm 0.2$ | $28.6 \pm 0.7$ |
| 300 | $63.6 \pm 0.2$ | $29.7 \pm 0.8$ |

An increase in PEG10k concentration from 0 to 300 mg/ml yields an ≈9 °C increase in the stability of G4 (Table 1). This is consistent with prior investigations of unlabeled G4 that found macromolecular crowding induced by PEG results in thermal stabilization by the reduction of water activity.^36^ The nuclear melting temperature is ≈14 °C lower than the melting temperature observed in vitro at near physiological crowding concentrations, 200 mg/ml PEG10k. Therefore, macromolecular crowding must not be the only effect (or dominating effect) modulating the stability of G4 inside cells. We hypothesize that chemical interactions must offset the stabilizing effect of macromolecular crowding on G4.

Macromolecular crowding alone also cannot explain the ≈19°C increase in iM stability observed in the nucleus of cells. Whereas a PEG titration from 0 to 300 mg/ml yields an ≈9°C increase in the stability of G4, it induces only an ≈2 °C stabilization of iM (Table 1). Our results match previous work on the c-MYC i-motif which found that 200 mg/ml PEG with sizes between 200 and 4000 kD induce only a 1-2 °C stabilization at neutral pH.^10^ That study showed that the same concentrations of PEG are more stabilizing at slightly-acidic, non-physiological pHs that promote hemi-protonation of the cytosine tracts. i-motifs are stabilized by base pairing between two cytosines, and optimal stability is achieved when one of the cytosines is protonated enabling three hydrogen bonds to form between the pair (Figure 1B).^37^ Thus, at physiological pH, the presence of PEG only modestly increases the stability of iM compared to dilute buffer conditions (Table 1, Figure S4B). While steric crowding effects likely contribute to the stabilization of iM inside the nucleus, there must be other interactions that contribute to the large ≈19 °C stabilization observed inside cells.

### Non-specific chemical interactions can be stabilizing or destabilizing, but no single effect replicates the nuclear stability trends of both G4 and iM

To isolate the effects of non-specific chemical interactions on G4 and iM, we measured their stabilities in vitro in buffers that mimic different cellular chemical interactions. We consider the effects of factors including the ionic strength, chemical composition of the nucleus, and the coexistence of nuclear DNA and proteins (Table S2, Table S4). All in vitro measurements were performed in 10 mM potassium phosphate, 200 mM KCl, 1 mM K2EDTA, pH 7.0 to closely mimic the ionic strength and ionic composition of cells.^38^

Whereas macromolecular crowding by PEG (purple) induces compaction of both DNAs compared to phosphate buffer alone (blue), chemical interactions have differential effects on G4 and iM (Figure 3). Non-specific chemical interactions in the nuclei are mimicked in vitro by small molecules, NER (green), and dilute solutions of macromolecules, ssDNA (orange) and BSA (magenta). The D/A at the beginning and end of the thermal melt reports on the end-to-end distance and structure of the folded and unfolded state, respectively. Although PEG has a larger effect on iM than G4, in both cases the initial and final D/A values are smaller (Figure 3), indicating that in crowded conditions the DNA accesses more compacted folded and unfolded states. Mimics of chemical composition of the nucleus on G4 are all destabilizing and result in slightly expanded folded states of G4, with more variability in the effect on the unfolded state (Figure 3A). Similarly to G4, small molecules have the smallest effect on the structure of iM (Figure 3, green). Conversely from G4, chemical interactions with macromolecules promote more compacted folded and slightly expanded unfolded states of iM (Figure 3B). Although a melting temperature can be extracted from iM measurements made in potassium phosphate alone (Figure 3B blue, Figure S4B), the small ΔD/A value with increasing temperature supports CD measurements that show iM is predominantly disordered (Figure S1C). Thus, chemical interactions are an important regulator of G4 and iM structure and stability.

**Figure 3:**
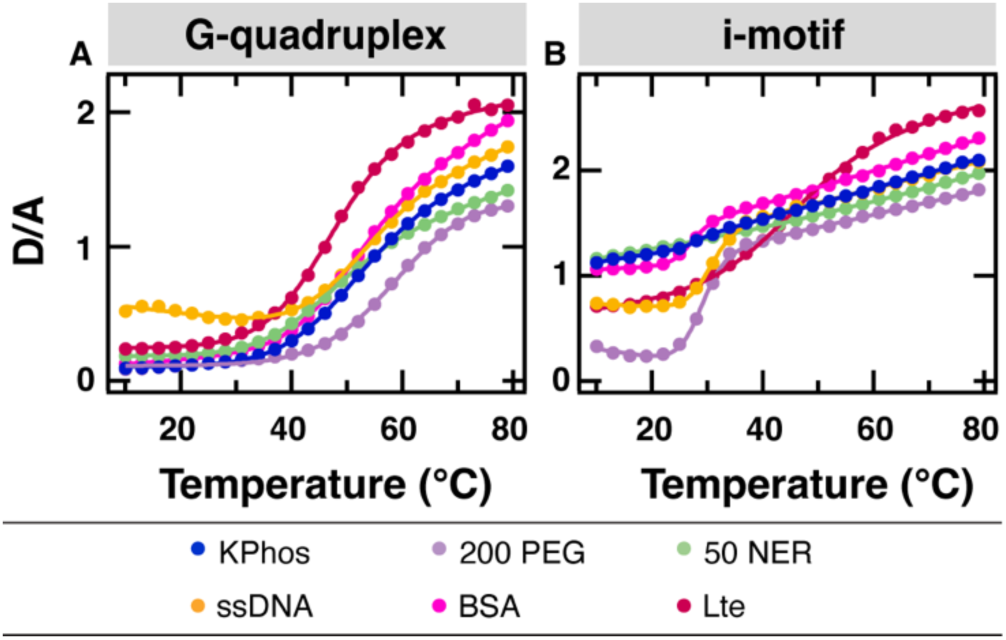
Chemical interactions are important regulator of G4 and iM structure and stability. Representative equilibrium temperature denaturation of **(A)** G4 and **(B)** iM monitored by fluorescence spectroscopy excited at 480 nm and collected from 475 to 700 nm. The signal from two-color FRET experiments is reported as a D/A ratio. Experiments were conducted in 10 mM potassium phosphate, 200 mM KCl, 1 mM K2EDTA, pH 7.0 alone (blue) or with 200 mg/ml PEG (purple), 50% NER (green), 10 mg/ml ssDNA (yellow), or 0.5-1 mg/ml nuclear cell lysate (red).

Lysis buffers have been engineered to maintain the structure and activity of proteins, and therefore likely replicate the effects of many non-specific interactions present in cells.

Cytosolic and nuclear lysis buffers have been previously used to mimic influences from non-specific interactions in their respective compartments.^22,39,40^ In this spirit, we performed experiments in nuclear lysis buffer (NER, ThermoFisher Scientific) to mimic non-specific interactions with ions (≈200 mM NaCl), small organic molecules (PBS, ≈5 mM EDTA), and short-chain fatty acids (≈0.1% Triton X-100) within the nuclei of cells (Patent No. 9580462 B2). We observed a linear trend in destabilization of G4 with increasing concentrations of NER, with 50 %v/v best matching the stability inside the nuclei of U-2 OS cells (Figure 4, Figure S5A). However, NER could not be used to replicate the in-cell stability of iM. Similar to G4, iM is destabilized by NER; At %v/v >50% the i-motif was so destabilized that the melting temperature could not be confidently measured (Figure 4, Figure S5B-C). That the stability of iM in NER is opposite of the nuclear stability suggests that the difference between nuclear and in vitro stability is not due to non-specific interactions with small organic molecules and metabolites.

**Figure 4:**
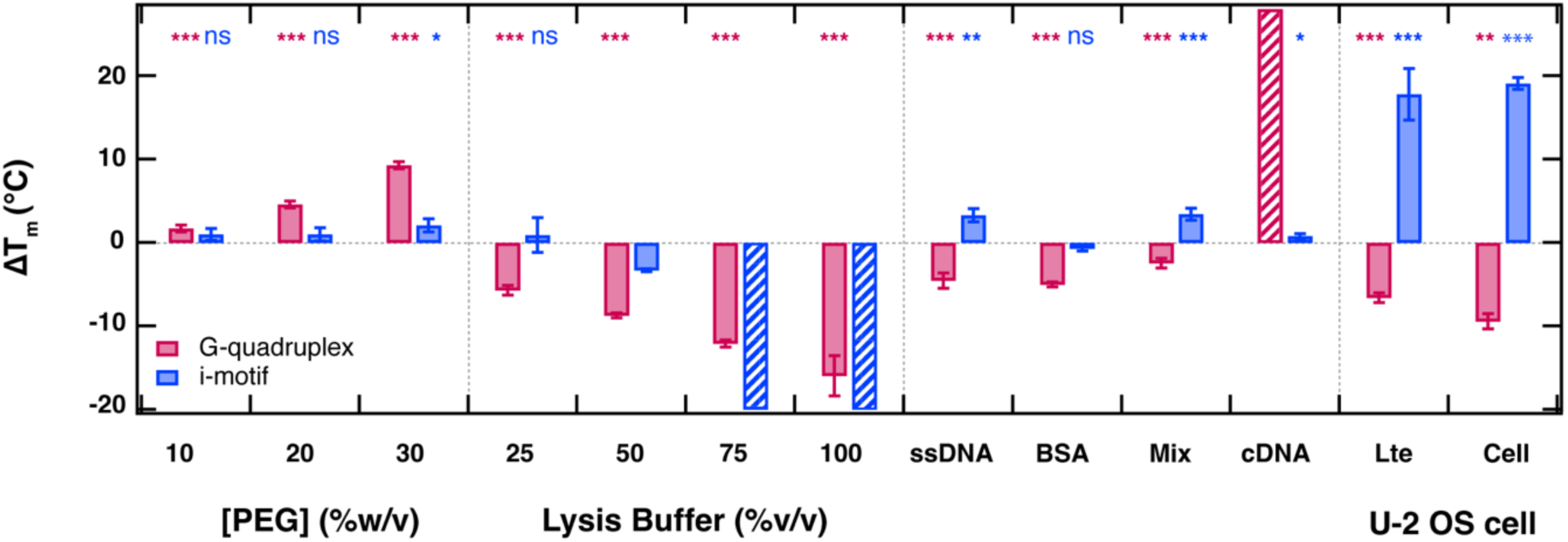
Measurements of G4 and iM stability in nuclear cell lysate most closely mimic the stability inside the nucleus. Effect of macromolecular crowding and chemical interactions on the stability of G4 (magenta) and iM (blue). To visualize stability differences, we plotted the change in melting temperature (ΔTm), calculated as the average melting temperature of each test condition minus that of the standard buffer (10 mM potassium phosphate, 200 mM KCl, 1 mM K₂EDTA, pH 7.0). Test conditions included macromolecular crowding alone (10-30 %w/v PEG10k), chemical interactions with small molecules or macromolecules alone [25– 100 %v/v NER lysis buffer; 10 mg/ml salmon sperm DNA (ssDNA); 0.5 μM complementary DNA (cDNA); 1 mg/mL BSA], and a mixture of the two [15% w/v PEG10k, 37.5 %v/v NER, 2.5 mg/ml ssDNA, and 10 mg/ml BSA in standard buffer (Mix)]. Also shown are stability differences between standard buffer and U-2 OS nuclear lysate (Lte), and between standard buffer and the nucleus of a living U-2 OS cell (Cell). Bars filled with diagonal stripes are conditions where the stability was too high or low to be confidently fit. Error bars are derived from propagating the standard error of three experiments in phosphate buffer and the standard error of three experiments for each respective condition (one experiment for iM in 50% NER, ≈40 measurements for G-quadruplex, and ≈60 measurements for i-motif in the nucleus of U-2 OS cells). \*\*\**p*<0.001; ** *p*<0.01; \**p*<0.05; ns *p>*0.05 between control condition (standard buffer) and test condition by a two-tailed *t* test. Statistical tests were not performed for conditions where the T_m_ was not quantified or only one measurement is reported.

Repulsions arising from the negatively charged phosphate backbone of DNA are an important factor in the electrostatic interactions that govern the thermodynamic stability of DNA.^41^ The repulsive strength of DNA in the nucleus has been found to stabilize RNA secondary structures and, therefore, likely also stabilizes DNA secondary structures like G4 and iM.^22,42^ To test this, we quantified G4 and iM folding in the presence of 10 mg/ml salmon sperm DNA (ssDNA), a mimic of non-specific electrostatic repulsion with nearby DNA molecules. In ssDNA, G4 is destabilized by ≈5 °C and iM is stabilized by ≈3 °C to 49.8 ± 0.9 °C and 30.8 ± 0.8 °C, respectively (Figure 4, Figure S6, Table S2, Table S4). Since the large opposing stabilities are not reproduced, non-specific electrostatic interactions with nucleic acids cannot completely explain the opposite stability trends of G4 and iM observed inside cells.

Another source of charge-charge interactions in the nucleus is proteins. Therefore, we also tested the effect of non-specific chemical interactions from proteins on the stability of G4 and iM using BSA since it is often employed in biochemical assays and is highly stable.^43^ While BSA destabilized G4 by ≈6 °C, the effect on iM was negligible (Figure 4, Tables S3 and

S5). Since iM is stabilized by ≈20 °C inside cells, non-specific interactions from proteins alone cannot justify the opposite trends.

Perhaps unsurprisingly, no single macromolecular crowding or non-specific chemical effect accounts for the nuclear stabilities of both G4 and iM. Rather, the different effects likely counteract each other to produce the observed in-cell stabilities. It is well established that the effects of steric crowding and non-steric chemical interactions are nonadditive.^22,40,44^ A true nuclear mimetic would produce all of these effects in tandem. To account for crowding and varying non-specific chemical effects, we created a mixture with 150 mg/ml PEG10K, 37.5 %v/v NER, 2.5 mg/ml ssDNA, and 10 mg/ml BSA. This results in an ≈3 °C destabilization of G4 and an ≈3 °C stabilization of iM (Figure 4, Figure S7). Evidently, the combination of crowding, ionic strength, electrostatics, and non-specific interactions with macromolecules reproduces the opposite trends but not the absolute magnitudes of G4 and iM stabilities observed in living cells. Thus, we hypothesize that specific interactions with macromolecules in the nucleus must drive the nuclear stabilities of G4 and iM.

### Weak, specific chemical interactions with nuclear proteins best reproduce the opposite nuclear stabilities of G4 and iM

First, we consider specific interactions between DNA; under native conditions the G- and C-rich strands of G4 and iM are present along with their complementary C- and G-rich strands, respectively. Therefore, we measured the stability of G4 and iM in vitro in the presence of their complementary DNA (cDNA) strands in a 1:1 ratio at neutral pH. Although iM is mildly stabilized, G4 exhibits large D/A values and a decrease in D/A with temperature consistent with the melting of duplex structure (Figure 4, Figure S8).^45^ This is consistent with previous fluorescence studies that showed at neutral pH, duplex structures largely outcompete formation of single-stranded G-quadruplex and i-motif structures.^45,46^ Since G4 and iM structures cannot compete with duplex formation in cells, cDNA is likely not the key contributor to the opposing nuclear stability trends.

Another possibility is that weak, specific interactions with other macromolecules in the nucleus are the major modulators of G4 and iM stability. Recently, weak interactions from proteins in the nucleus were shown to have a direct effect on DNA G-quadruplexes inside cells.^47^ Therefore, we used dilute U-2 OS nuclear cell lysate (Lte) to mimic the contributions from specific interactions with nuclear proteins to DNA folding. Initial D/A values show that Lte promotes folding of both DNAs (Figure 3, red). Intriguingly, both DNAs access more extended unfolded states than in any other condition tested, suggesting that interactions with nuclear proteins promote more extended unfolded states. The stability trends in Lte most closely resemble in-cell data for both G4 and iM (Figure 4, Figure S9). Previous protein folding and binding studies have identified cell lysate as the most accurate mimic for in-cell dynamics.^23,44^ Because the concentration of proteins in the nuclear cell lysate is a fraction of what is present inside cells, our measurements can reproduce the opposite stability trends but not the absolute melting temperatures of G4 and iM in living U-2 OS cells. Nonetheless, our experiments in nuclear cell lysate show that specific interactions with proteins within the nucleus are the major contributors to the stability of G4 and iM inside cells.

### Nuclear proteins cannot account for in-cell folding dynamics

In addition to thermodynamics, we compared the folding dynamics of G4 and iM in vitro and in cells. To extract relaxation lifetimes, we monitored the D/A signal over time for a temperature jump to T_m_. The transients were fit to a double exponential function where the first exponential accounts for the instrument response and unresolved dynamics and the second exponential are the DNA dynamics (Equation 2). Our fit includes a stretch factor that provides information about deviations from two-state behavior.

In aqueous buffer, G4 has a relaxation lifetime, τ, of 0.19 ± 0.03 s (n=16) which is in good agreement with previous work.^48^ However, folding inside cells is an order of magnitude slower, 7 ± 1 s (n=17) (Figure 5A). Because G4 was slow to reach equilibrium inside cells, we extended the length of our jumps from 5 seconds to 20 seconds (Figure S10). No significant deviations from two-state behavior were found for measurements in aqueous buffer, which required a β value of 1 to be fitted with high confidence. While the average β value in cells was 1.0 ± 0.1, β ranged from 0.72 to 2.3 deviating far from a single exponential function. This can occur when there are multiple populations with different relaxation lifetimes that are averaged together in an ensemble measurement. β values > 1 may also indicate that the folding mechanism is multi-state inside the cell. Indeed, it is well established that the folding landscape of TBA is more rugged and complex than that of a simple two-state model;^49^ in vitro, the transitions between local intermediate states occur on the sub-millisecond timescale and cannot be captured by FReI, likely giving rise to the apparent two-state behavior. Our data suggest that the folding of intermediate states is slowed inside cells, that new slow intermediates states are accessed, and/or that there are multiple populations with different relaxation lifetimes.

**Figure 5:**
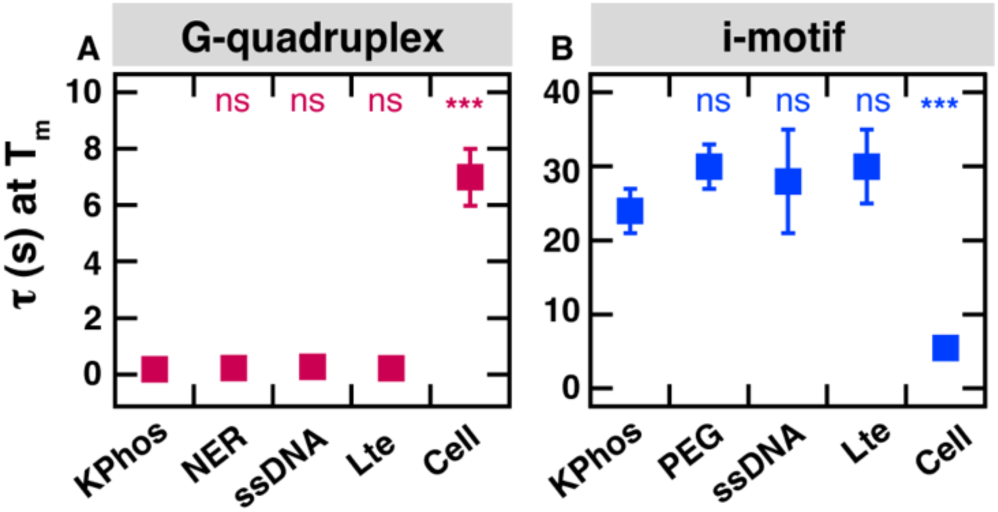
G4 folding is slow and iM folding is fast inside cells compared to in vitro conditions. Average relaxation lifetimes (τ) of **(A)** G4 and **(B)** iM following a temperature-jump to T_m_ in KPhos (10 mM potassium phosphate, 200 mM KCl, 1 mM K2EDTA, pH 7.0), 200 mg/ml PEG, 50 %v/v lysis buffer (NER), 10 mg/ml ssDNA, dilute nuclear cell lysate (Lte), or the nuclei of fifteen U-2 OS cells. Data was fitted to Equation 2. Data could not be fit for G4 in PEG (too fast) and iM in NER lysis buffer (too small amplitude). Error bars indicate standard error of the mean of three measurements for in vitro conditions and approximately fifteen measurements for cells. \*\*\**p*<0.001 and ns *p*>0.05 between experiments (n=3) in standard buffer (KPhos) and each respective condition derived with a two-tailed *t* test.

While G4 folding is slowed down inside cells, the opposite was observed for iM. Relaxation lifetimes of iM were four times faster inside cells compared to in vitro (Figure 5B). In vitro, the dynamics were so slow that iM required longer t-jumps to achieve equilibrium. Therefore, the length of t-jumps was extended from 10 seconds to 90 seconds to accommodate the slower relaxation lifetimes (Figure S11). Such slow rates have been observed for iM where folding times can range from several minutes to hours.^50^ In aqueous buffer at neutral pH, the observed relaxation time constant is 24 ± 2 s with a β value of 1.5 ± 0.3. In cells, iM had an average relaxation lifetime of 5.5 ± 0.7 s (n = 14) with a β factor of 3.0 ± 0.5. The in-cell β value ranged from 1 to 7 (Table S3). β values > 1 are consistent with a multi-state mechanism but may also arise from heterogeneity in the population. Indeed, like G-quadruplexes, i-motif DNA formation is driven by kinetic partitioning, with the process involving multiple intermediate species.^27,50^

To test whether macromolecular crowding or chemical interactions can account for the folding kinetics of G4 and iM inside cells, we performed temperature jumps in PEG, NER, ssDNA, and Lte (Figure 5, Table S5-6). However, none of the relaxation lifetimes measured in these conditions vary significantly from standard phosphate buffer. Despite the good agreement between stabilities in cells and in nuclear cell lysates, the in-cell kinetics of G4/iM are not reproduced in nuclear cell lysate.

### Organization by phase separation is the key contributor to opposing G4 and iM kinetics

Prior studies show that G-quadruplexes and i-motifs localize in puncta in the nuclei of living cells.^12,13^ Indeed, we observe puncta in cells microinjected with G4/iM (Figure 2 insets). To examine the impact of phase separation on G4 and iM stability and kinetics, we induced phase separation in vitro with 0.1 mg/ml poly-L-lysine (PLL). PLL is a cationic homopolypeptide that promotes phase separation of negatively charged polymers like DNA. A turbidity assay revealed that phase separation occurs with concentrations as low as 5 μM for both unlabeled G4 and iM (Figure S12).

To test whether DNA colocalized with PLL into the droplets, 5 μM of Cy5.5-labeled G4/iM samples were imaged with 0.1 mg/ml FITC-labelled PLL. CD measurements confirmed that conjugation of the Cy5.5 fluorophore did not disrupt the formation of the G-quadruplex and i-motif structures (Figure S13). Upon incubation in FITC-labelled PLL, fluorescence imaging confirmed that both G4 and iM undergo phase separation into droplets (Figure 6).

**Figure 6:**
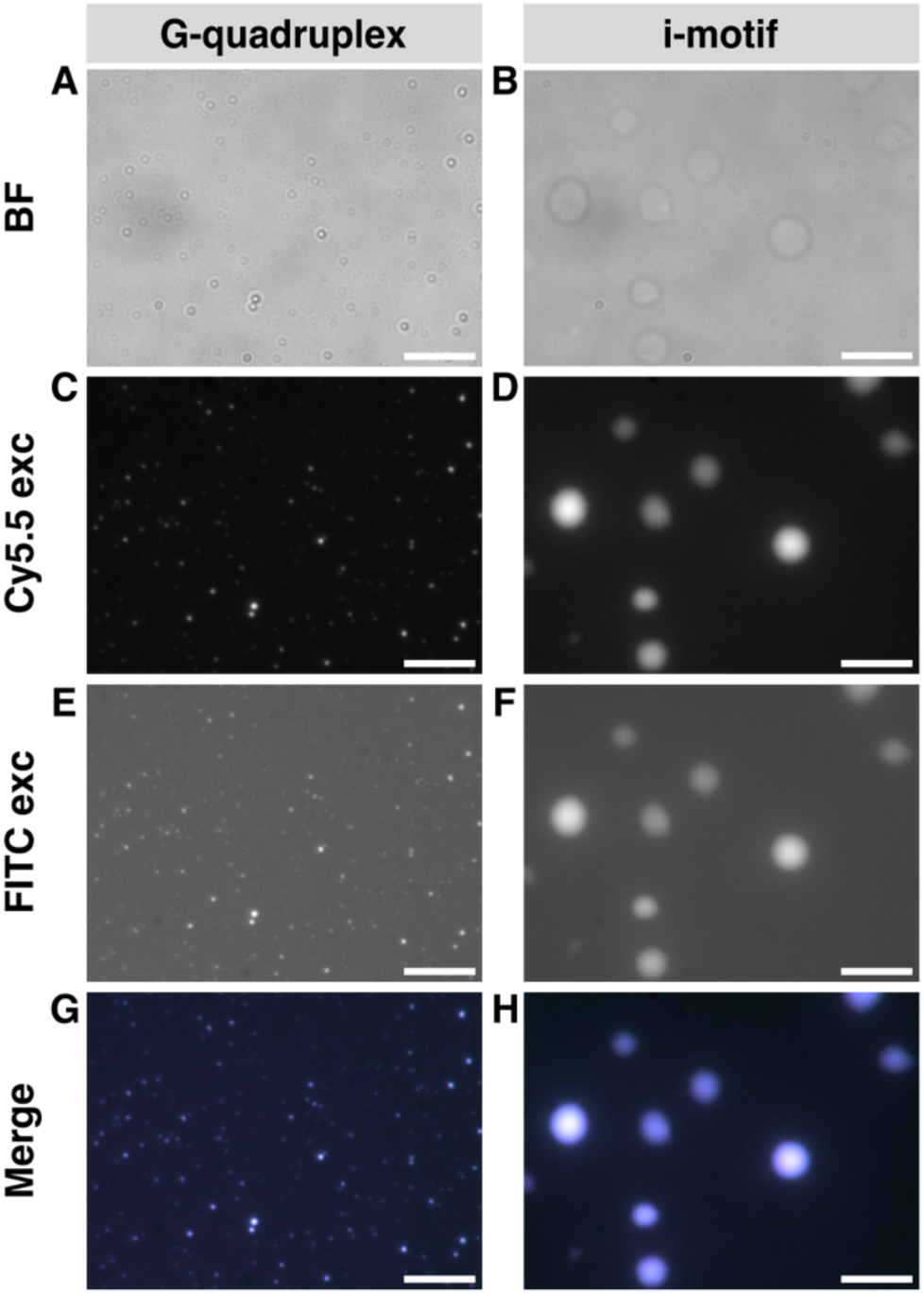
Fluorescence microscopy images show G4/iM and PLL colocalize into droplets in vitro. Brightfield images of 5 μM Cy5.5-labeled **(A)** G4 and **(B)** iM in the presence of 0.1 mg/ml FITC-labelled PLL in phosphate buffer (10 mM potassium phosphate, 200 mM KCl, 1 mM K_2_EDTA pH 7.0). Direct excitation of the red channel (**C-D**), green channel (**E-F**), and an overlay of both channels (**G-H**) show colocalization of Cy5.5-labeled G4/iM and FITC-labelled PLL into droplets. Scale bar is 5 μm.

Consistent with prior studies, iM-Cy5.5 droplets are larger than G4-Cy5.5 droplets due to the larger subunit size (Figures 6C-D).^51^ An overlay of the red and green channels confirmed that both Cy5.5-labeled DNA and FITC-PLL colocalized into the droplets (Figures 6G-H). Since an excess of FITC-PLL was added to the sample, much of the polymer appeared to be free in solution (Figures 6E-F). Intriguingly, more of the G4-Cy5.5 and FITC-PLL appear to be in the dense phase than for iM-Cy5.5 (Figure S14A). We therefore calculated the partitioning coefficient (*K_p_*) by comparing the fluorescence intensities of the DNA and PLL in the condensates to the surrounding dilute phase. The partitioning analysis can be used to compare the degree of condensation for the structures relative to the dilute phase.^52^ Although condensates contained more G4-Cy5.5 than iM-Cy5.5, the amount of FITC-PLL in the G4-Cy5.5 condensates was lower compared to those containing iM-Cy5.5 (Figure S14B). This suggests that the degree of interactions with proteins and their relative concentrations in condensates may be important in tuning the stability and folding of the structures.

We next examined the role of phase separation on the differential stability and kinetic trends of the FRET-labelled structures with temperature jumps on the PLL-induced droplets. In 0.1 mg/ml unlabelled PLL, both FRET-labelled G4 and iM localized into droplets (Figures 7A-B); G4 was destabilized, and the relaxation lifetime of G4 was slowed to least 20 s (Figure 7C, Table S2). Conversely, iM was stabilized, and the relaxation lifetime was sped to an average of 5 ± 2 s (Figure 6D, Table S4). Thus, measurements in condensates match the opposite kinetic trends of both G4 and iM inside living cells. Taken together, these results suggest that the folding of G4 and iM is tuned by associative phase separation with positively charged species in the nucleus.

**Figure 7:**
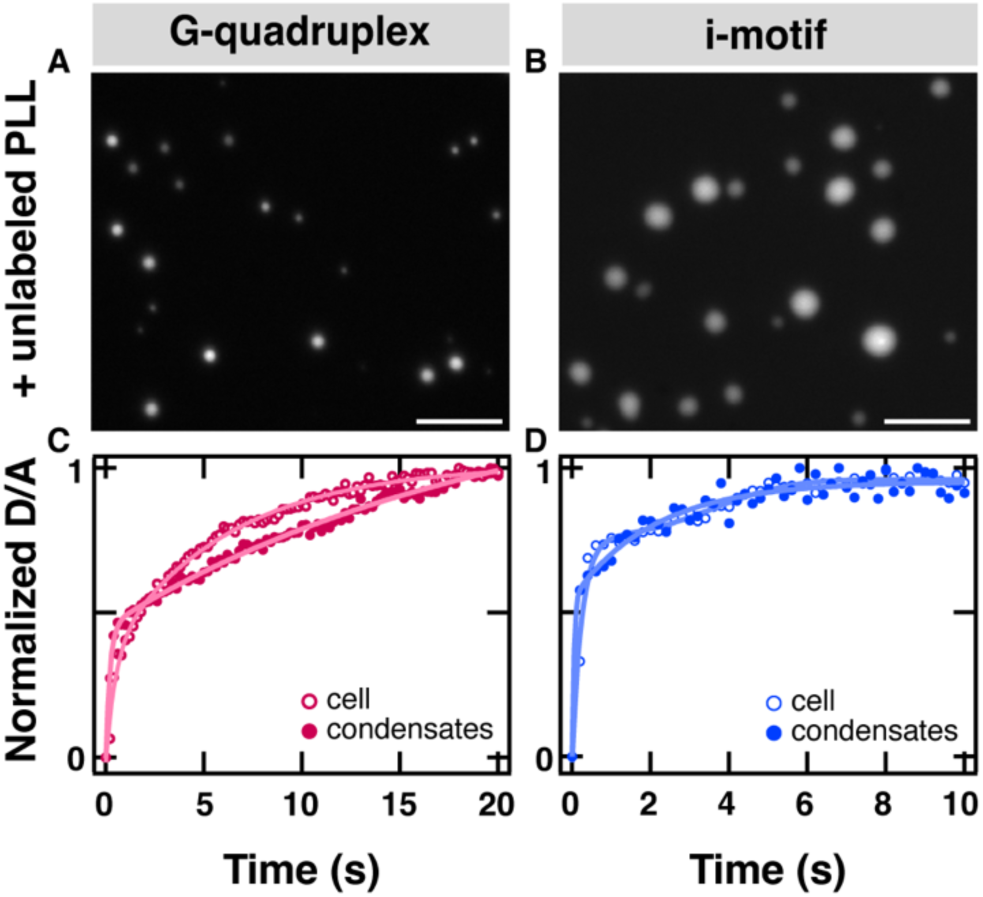
Phase separation of G4/iM in the presence of positively charged species most closely mimics in-cell kinetics. Images of 5 μM FRET-labelled **(A)** G4 and **(B)** iM in the presence of 0.1 mg/ml unlabelled poly-L-lysine (PLL) in phosphate buffer (10 mM potassium phosphate, 200 mM KCl, 1 mM K_2_EDTA pH 7.0). Scale bar is 5 μm. Relaxation lifetimes extracted from the condensates containing FRET-labelled **(C)** G4 and **(D)** iM containing 0.1 mg/ml unlabelled PLL in phosphate buffer at pH 7.0.

## DISCUSSION

The model TBA G-quadruplex, G4, is destabilized by ≈1 kcal/mol inside cells compared to in vitro. This degree of destabilization may be sufficient to abrogate persistent G-quadruplex formation that would otherwise lead to disruption in active replication during the S phase where G-quadruplexes are particularly enriched. Indeed, recent cryo-EM studies revealed G-quadruplex-induced stalling of replisome progression due to lodging of the G-quadruplex structure within the CMG helicase.^53^ The study highlighted that rapid turnover of the G-quadruplex structure is crucial to prevent replisome stalling and maintain genome stability. Yet we also observed significantly slower folding of G4 inside cells compared to dilute conditions in vitro. Phase separation facilitated by charge-driven interactions with IDRs of proteins can lead to condensation of G-quadruplexes.^54–57^ Such G-quadruplex-protein interactions are reported to be on the seconds to minutes timescales in vitro,^58^ consistent with our in-cell and in vitro condensate measurements. Thus, it is likely that phase separation, facilitated via interactions with such proteins, orchestrates a nuclear environment that dynamically modulates G4 folding, accessibility, and turnover.

The model human telomeric i-motif, iM, is stabilized by ≈1.2 kcal/mol inside cells compared to in vitro crowding conditions (200 mg/ml PEG). Indeed, prior studies show that i-motif sequences can form stable structures inside cells, although their propensity to form is dependent on both sequence and the cell cycle.^13,14,59^ It has been observed that various i-motif sequences can interact with endogenous proteins. Our results align with these findings and also demonstrate that such interactions stabilize iM. Additionally, we observed that iM kinetics were 16x faster inside cells. While the kinetics of i-motif structures inside condensates are largely unexplored, i-motif puncta inside cells are known to exhibit dynamic assembly during varying stages of the cell cycle and kinetically-driven species are largely more favorable.^14,50^

Contrasting folding behavior of iM and G4 inside cells, which can only be reproduced by phase separation, highlights the central role of condensation in the stability and dynamics of the structures. Our results support a model whereby interactions with nuclear proteins and associative phase separation provide exquisite control over the stability and kinetics of non-canonical DNA structures inside cells. Numerous studies have shown that interactions with proteins and phase separation can be either stabilizing or destabilizing for various G-quadruplex and i-motif sequences.^27,55,60,61^ However, independent of the sequence, we propose that specific interactions are necessary to tune the folding behavior to biologically relevant timescales. Past work, combined with our own quantitative comparison between in vitro and in cell, therefore implies that interactions with proteins can tune the stability of these non-canonical structures (either stabilizing or destabilizing) to promote their role as regulatory elements in active replication and transcription. Together, our results shed light on how the intranuclear environment modulates the assembly of these structures and underscores the importance of studying DNA stability and dynamics inside cells. This is especially crucial for biomedical applications where the effectiveness of therapies relies on targeting the correct ensemble of states.

## ACKNOWLEDGEMENTS

This work was supported by National Institutes of Health (NIH) grant R35 GM151146. B.P. was partially supported by the NIH under Chemical Biology Training Grant T32 GM067543.

C.H. was partially supported by the Yale College Dean’s Office through the STARS II Program. This research made use of the Chemical and Biophysical Instrumentation Center at Yale University (RRID:SCR_021738).

## AUTHOR CONTRIBUTIONS

B.P., H.Y. and C.M.D. conceived the study. B.P. and H.Y. carried out the investigation. B.P., C.H. and C.M.D performed formal analysis. B.P. wrote the initial draft. B.P. and C.M.D. wrote the manuscript and generated the visualizations therein. C.M.D. provided supervision, project administration, and funding.

## SUPPLEMENTARY DATA

Supplementary Data are available at NAR online.

## DATA AVAILABILITY

All data are available in the article or Supplementary Information. Raw data will be shared upon request to the corresponding/first author.

