## Supporting Information for "Phase Separation Tunes the Stability and Dynamics of G-quadruplex and i-Motif DNA in the Nuclei of Living Cells"

[2] Present Address: NYU Langone Health, New York, New York, 10016, United States

[3] Present Address: Physical Sciences Inc., Andover, Massachusetts, 01810, United States

### METHODS

**Circular Dichroism (CD) Spectroscopy.** CD spectra were recorded on a Chirascan spectrophotometer (Applied Photophysics, Charlotte, NC) connected to a Quantum Northwest TC125 temperature control module. All measurements were obtained using a 2 mm pathlength cuvette (Starna Cells, Atascadero, CA). Wavelength scans were recorded over the range of 340 to 210 nm in 1 nm intervals and a bandwidth of 1 nm, and a time-per-point of 0.5 sec was used for spectral acquisition. Thermal unfolding experiments were performed with a temperature range of 10 to 88 °C using a ramp rate of 0.5 °C/min. During the thermal unfolding experiment a full wavelength scan was obtained in 3 °C intervals after a 180 sec delay. The sample was covered with mineral oil to prevent evaporation during thermal melts. Data is reported as the difference in molar extinction coefficients to normalize for nucleic acid concentration and path length of the cuvette.[1]

### DISCUSSION

**Formation of FRET-labeled G4 and iM confirmed by circular dichroism**. G4 is an anti-parallel G-quadruplex with a characteristic CD spectrum with positive bands near 245 nm and 295 nm and a negative band near 265 nm.[2–4] Figure S1A shows that the CD spectrum for FRET-labeled G4 is in good agreement with the unlabeled construct, indicating little to no perturbations to its secondary structure by the addition of fluorophores. The stability of the labeled and unlabeled G4 was assessed by temperature-dependent CD measurements (Figure S1B). Comparison of the melting temperatures at 295 nm of the unlabeled G4 (54 ± 1) and FRET-labeled G4 (47 ± 1) shows a slight destabilization upon addition of the fluorophores. Nonetheless, the overall topology of the G-quadruplex is preserved in the slightly destabilized FRET-labeled construct.

CD measurements were also performed on unlabeled and FRET-labeled iM. In potassium phosphate buffer at neutral pH, the CD spectrum of FRET-labeled iM overlaps with the unlabeled sequence (Figure S1C), revealing minimal differences in secondary structure. The positive band at 276 nm and negative band at 247 nm are consistent with the CD spectrum of single stranded unordered C-rich DNA, suggesting that under these conditions the DNA is predominantly unfolded.[5] This is because at neutral pH, all the cytosines are deprotonated and are unable to form the $C{\cdot}C^{+}$ base pairs required for i-motif formation. Macromolecular crowding can promote the folding of i-motifs. Indeed, a partially folded i-motif structure is observed at neutral pH in the presence of 20 %w/v PEG10k (Figure S1C). This is evident with a red shift in the spectra, resulting in a positive band at 287 nm and a negative band at 253 nm. A completely folded i-motif structure has characteristic CD bands at 287 and 260 nm.[5,6] Therefore, we extracted the melting temperatures of unlabeled and FRET-labeled iM under molecular crowding conditions to quantify changes in the stability upon addition of the fluorophores. In the presence of 20 %w/v PEG10k, the melting temperatures of unlabeled and FRET-labeled iM are 30 ± 0.2 °C and 26 ± 0.1 °C, respectively (Figure S1D). Although we observe a decrease in the melting temperature by 4 °C upon labeling, the labeled construct is still capable of folding into an i-motif structure at near physiological conditions.

**FRET-labeling minimally perturbs G4 and iM structure and thermal stability.** The in vitro FRET detected melting temperatures are 3-4 °C higher than the melting temperatures extracted from CD measurements (Figure S1B, Figure S1D, Figure 2). Because CD spectroscopy monitors global changes in secondary structure whereas FRET probes the end-to-end distance between the two fluorophores at the termini of the DNA, such differences in melting temperatures are not surprising. Our results suggest that in both cases the DNA tertiary structure unfolds prior to the termini coming apart. Nonetheless, CD melting experiments confirm that conjugation of the dyes to the oligonucleotides does not disrupt their ability to fold into G-quadruplex and i-motif structures and demonstrate that FRET can report on their temperature-induced unfolding. Therefore, FRET-labeled G4 and iM will serve as a model for comparison of G-quadruplex and i-motif folding in vitro and in cell.

**Ionic strength influences the stability of G4 and iM**. Our standard buffer has 200 mM KCl because $K^+$ is the most common ion in the intracellular fluid of mammalian organisms and the net concentration of ions in a cell is ≈200 mM.[7] We also investigated the folding of G4 and iM in DPBS. DPBS was engineered to mimic the physiology of the intracellular fluid; the dominant ion in the intercellular fluid is $Na^+$ and the net concentration of ions is ≈150 mM.[7] Formation of G-quadruplexes is facilitated by the presence of several cations including sodium and potassium, and G-quadruplexes are known to be particularly stabilized by potassium ions in vitro.[8] As expected, G4 is destabilized by ≈25 °C in DPBS (Figure S3A, Table S3). Of all chemical interactions tested this by far had the largest effect on G4; however, DPBS does not reproduce the intracellular physiology and, therefore, cannot explain the ≈10 °C destabilization of G4. DPBS also does not replicate the ≈19 °C in-cell stabilization of iM. Like G-quadruplexes, i-motifs are sensitive to ionic strength and identity.[9] The stability of iM is relatively unchanged in DPBS, but with a lower enthalpy change (Figure S3B, Table S5) indicating a less cooperative transition. Therefore, ionic strength cannot explain the differential stability trends observed inside cells and in vitro.

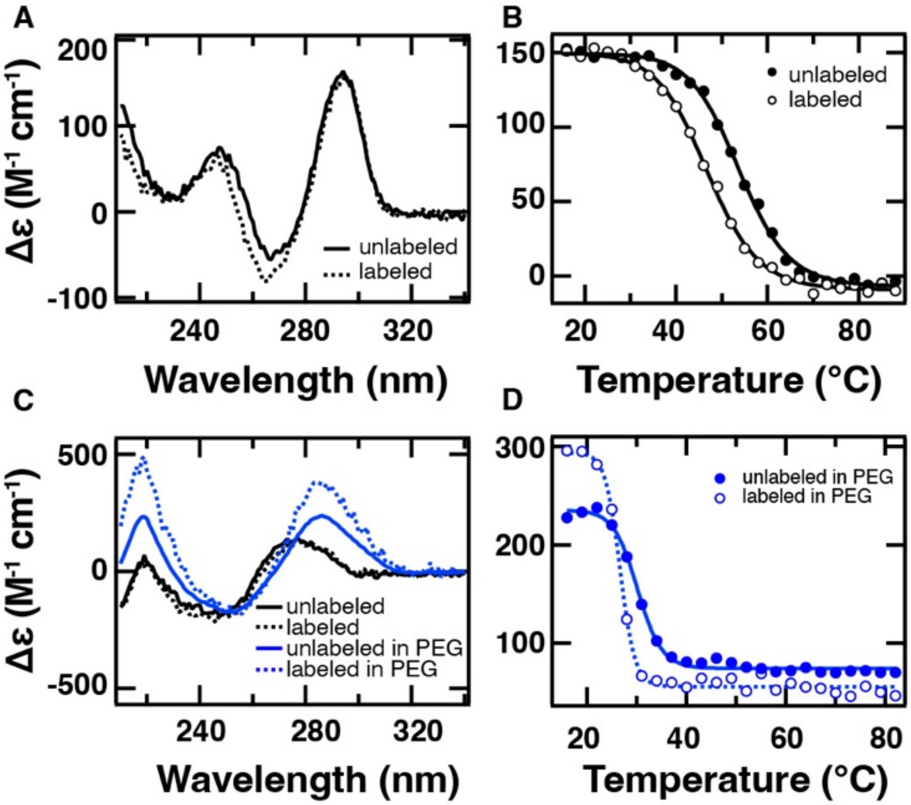


**Figure S1.** Representative far-UV CD spectra of 2 μM unlabeled (solid line) and FRET-labeled (dashed line) **(A)** G4 and **(C)** iM in 10 mM potassium phosphate, 200 mM KCl, 1 mM $K_2EDTA$, pH 7.0 alone (black) or with 200 mg/ml PEG10k (blue). Thermal denaturation of unlabeled (closed circles) and FRET-labeled (open circles) **(B)** G4 monitored by CD at 295 nm and **(D)** iM monitored by CD at 287 nm. Continuous line overlaid on thermal melts represent a fit to Equation 1.

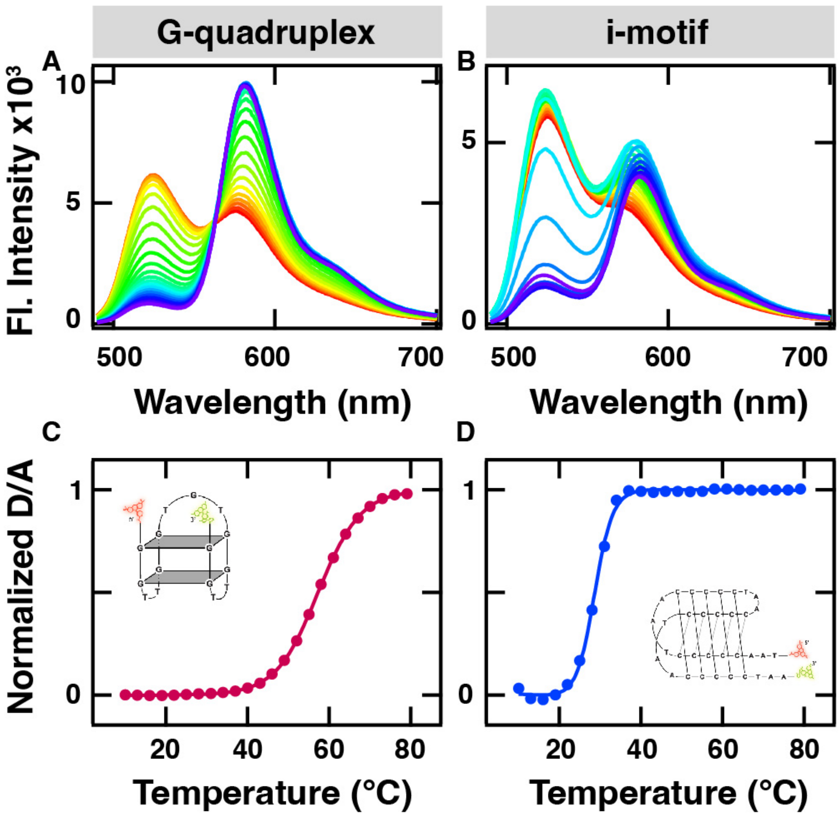


**Figure S2**. Representative temperature-dependent fluorescence emission spectra of 0.5 µM **(A)** G4 and **(B)** iM in 200 mg/ml PEG10k (10 mM potassium phosphate, 200 mM KCl, 1 mM $K_2$EDTA, pH 7.0) collected from 10 °C to 79 °C in 3 °C intervals. The FRET-labeled DNA were excited at 480 nm and monitored from 475 to 700 nm. (**C-D**) The donor/acceptor (D/A) ratio at each temperature was calculated from the donor (D) and acceptor (A) emission intensities at 520 nm and 580 nm, respectively. The temperature dependent D/A ratios are fit (continuous line) to a two-state equilibrium model (Equation 1) and normalized to correct for pre- and post- transition baselines.

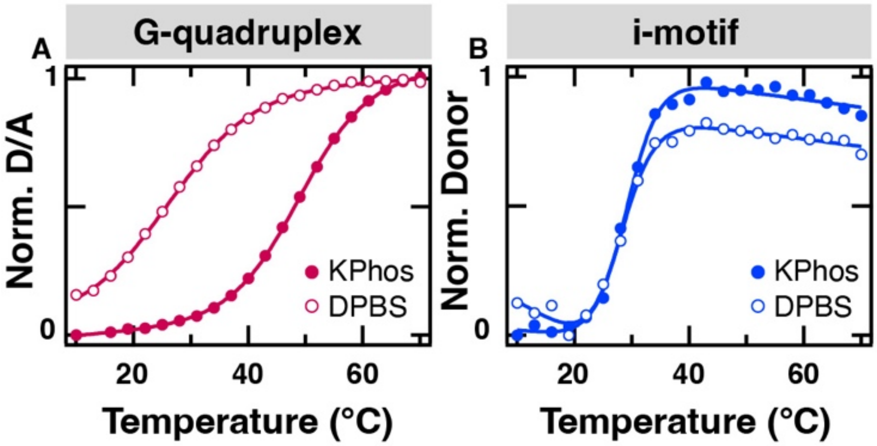


**Figure S3.** Representative thermal denaturation of 0.5 μM **(A)** G4 and **(B)** iM in 10 mM potassium phosphate, 200 mM KCl, 1 mM $K_2$EDTA, pH 7.0 (closed circles) and DPBS pH 7.0 (open circles) monitored by fluorescence spectroscopy. DNA were excited at 480 nm and monitored from 475 to 700 nm. Continuous line overlaid on thermal melts in represent a fit to Equation 1.

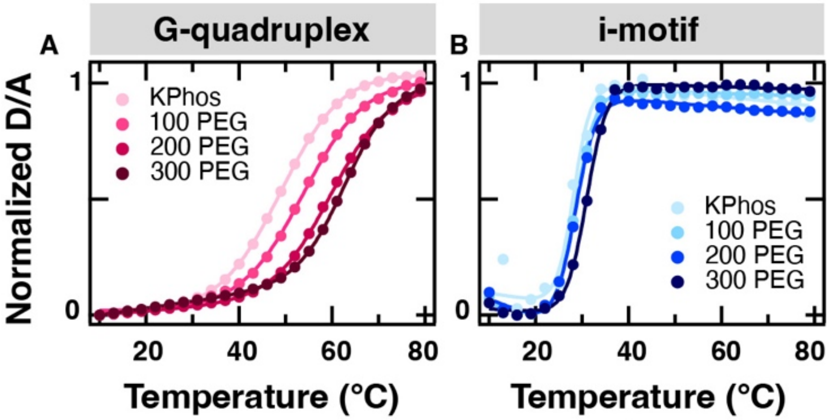


**Figure S4.** Representative thermal denaturation of 0.5 μM **(A)** G4 and **(B)** iM in increasing concentration of PEG (0, 100, 200, or 300 mg/ml PEG 10k in 10 mM potassium phosphate, 200 mM KCl, 1 mM $K_2$EDTA, pH 7.0). DNA samples were excited at 480 nm and monitored from 475 to 700 nm by fluorescence spectroscopy. Continuous line overlaid on thermal melts represent a fit to Equation 1.

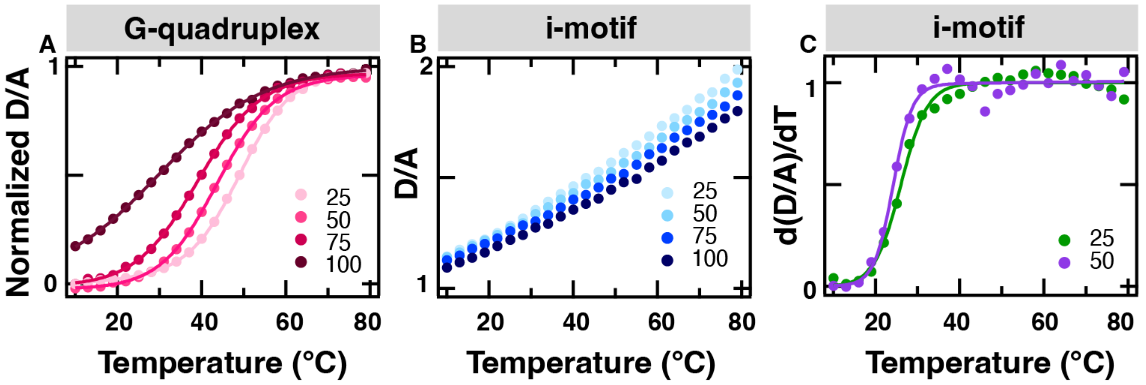


**Figure S5.** Representative thermal denaturation of 0.5 μM **(A)** G4 and **(B)** iM in increasing concentration (%v/v) of NER in standard buffer (10 mM potassium phosphate, 200 mM KCl, 1 mM $K_2$EDTA, pH 7.0) and (**C**) representative first derivative of iM in 25 (green) and 50 (purple) (%v/v) NER in standard buffer. DNA samples were excited at 480 nm and monitored from 475 to 700 nm by fluorescence spectroscopy. Continuous line overlaid on thermal melts in **A** and **C** represent a fit to Equation 1. The iM data in **B** could not be fit to Equation 1 and, therefore, is not normalized.

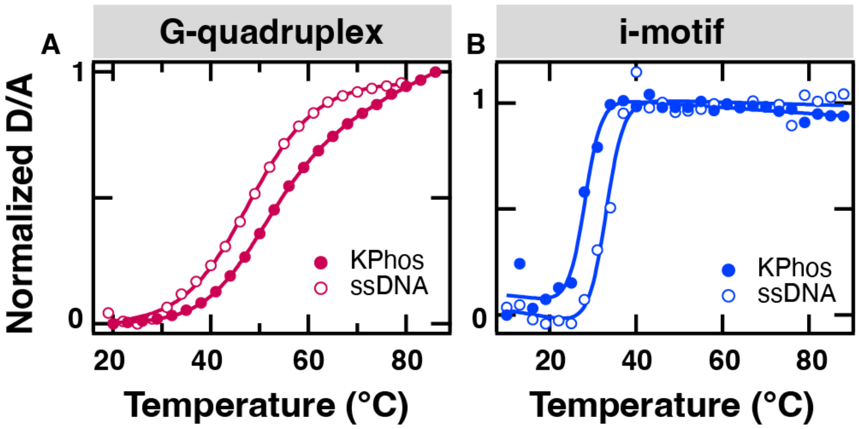


**Figure S6.** Representative thermal denaturation of 0.5 µM **(A)** G4 and **(B)** iM in 10 mM potassium phosphate, 200 mM KCl, 1 mM $K_2$EDTA, pH 7.0 with (open circles) and without (closed circles) 10 mg/ml salmon sperm DNA. DNA samples were excited at 480 nm and monitored from 475 to 700 nm by fluorescence spectroscopy. Continuous line overlaid on thermal melts in represent a fit to Equation 1.

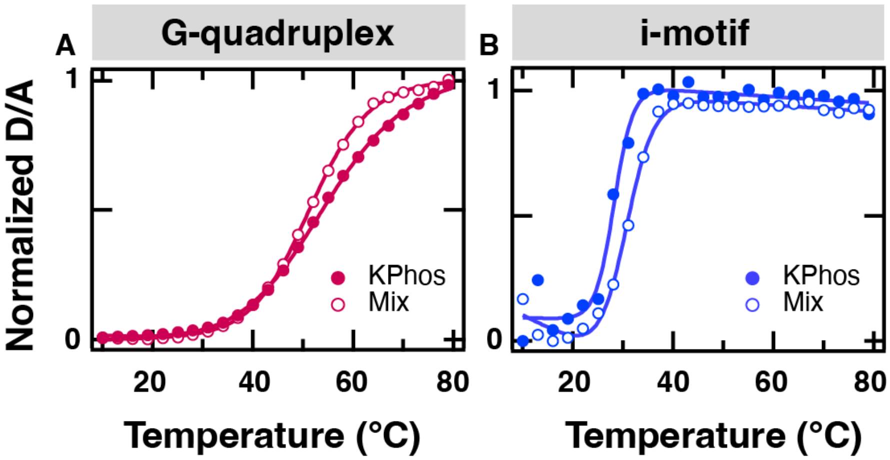


**Figure S7.** Representative thermal denaturation of 0.5 µM **(A)** G4 and **(B)** iM in 10 mM potassium phosphate, 200 mM KCl, 1 mM $K_2EDTA$, pH 7.0 with (open circles) a mixture containing 150 mg/ml PEG10k, 37.5 (%v/v) NER, 2.5 mg/ml ssDNA, and 10 mg/ml BSA or without (closed circles). DNA samples were excited at 480 nm and monitored from 475 to 700 nm by fluorescence spectroscopy. Continuous line overlaid on thermal melts represent a fit to Equation 1.

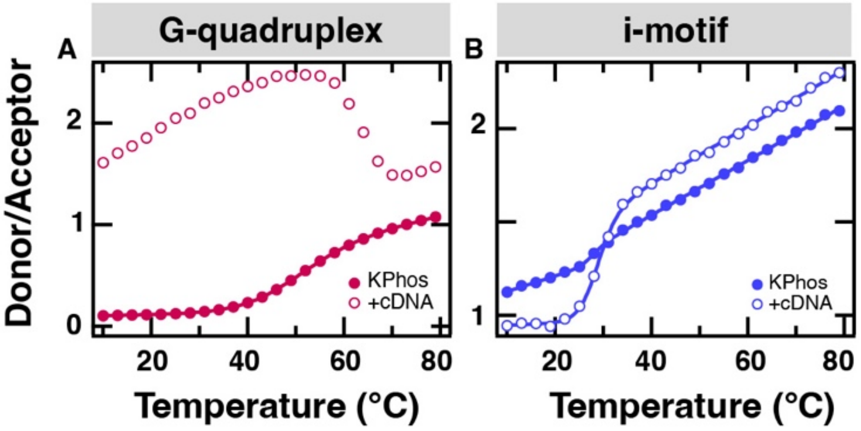


**Figure S8.** Representative thermal denaturation of 0.5 μM **(A)** G4 and **(B)** iM in the presence of complementary DNA at a 1:1 ratio in 10 mM potassium phosphate, 200 mM KCl, 1 mM $K_2EDTA$, pH 7.0. G4/iM and complementary DNA were annealed separately in the thermocycler and mixed before thermal melt. DNA samples were excited at 480 nm and monitored from 475 to 700 nm by fluorescence spectroscopy. Data collected from 10 °C to 79 °C in 3 °C intervals.

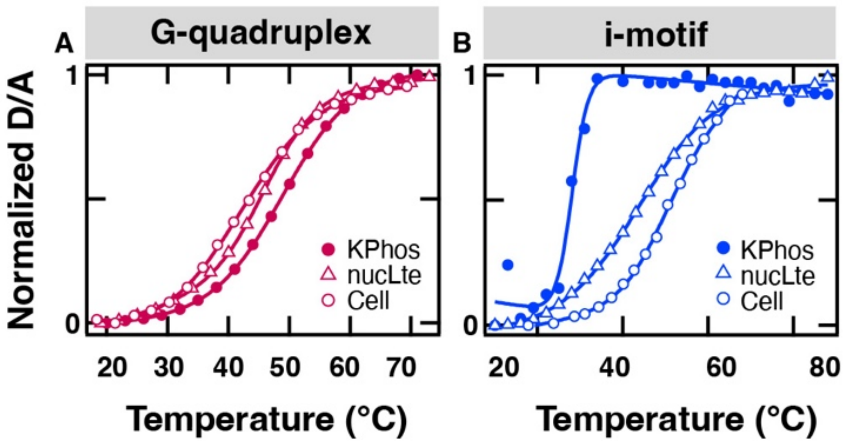


**Figure S9.** Representative thermal denaturation of 0.5 µM **(A)** G4 and **(B)** iM in 10 mM potassium phosphate, 200 mM KCl, 1 mM $K_2$EDTA, pH 7.0 (closed circles), in a U-2 OS cell (open circles), and in 0.5-1.5 mg/ml nuclear cell lysate dialyzed in in 10 mM potassium phosphate, 200 mM KCl, 1 mM $K_2$EDTA, pH 7.0 (triangles). Continuous line overlaid on thermal melts represent a fit to Equation 1.

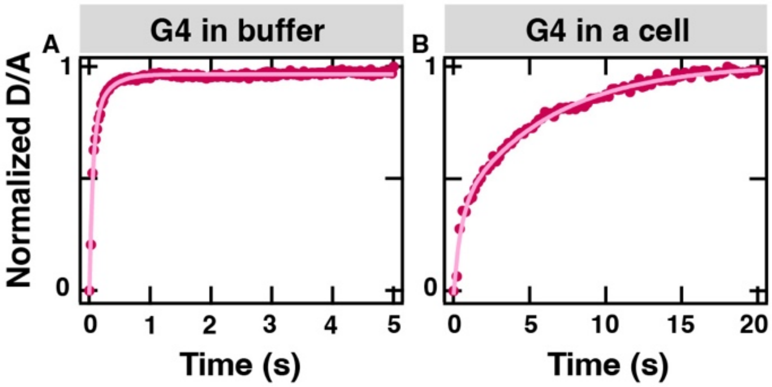


**Figure S10.** Normalized donor/acceptor kinetic trace of G4 in **(A)** potassium phosphate buffer (10 mM potassium phosphate, 200 mM KCl, 1 mM $K_2EDTA$, pH 7.0) (closed circles) and **(B)** a representative U-2 OS cell (closed circles) obtained by FReI microscopy. The kinetic traces are extracted for the temperature jump to the midpoint of unfolding. Continuous line overlaid on thermal melts represent a fit to Equation 2.

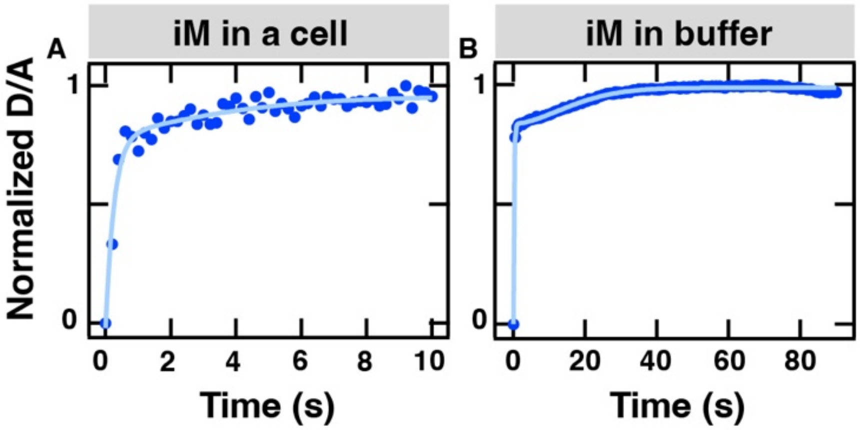


**Figure S11**. Normalized donor/acceptor kinetic trace of iM in **(A)** a representative U-2 OS cell (closed circles) and in **(B)** potassium phosphate buffer (10 mM potassium phosphate, 200 mM KCl, 1 mM $K_2$EDTA, pH 7.0) (closed circles) obtained by FReI microscopy. The kinetic traces are extracted for the temperature jump to the midpoint of unfolding. Continuous line overlaid on thermal melts represent a fit to Equation 2.

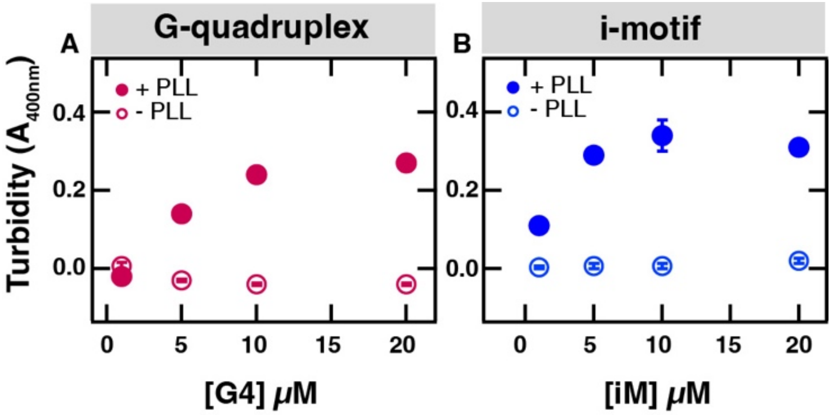


**Figure S12**. Turbidity measured by tracking absorbance at 400 nm of unlabeled **(A)** G4 or **(B)** iM in the presence (closed circles) or absence (open circles) of 0.1 mg/ml poly-L-lysine in KPhos buffer (10 mM potassium phosphate, 200 mM KCl, 1 mM $K_2$EDTA pH 7.0). Error bars indicate the standard deviation of three measurements.

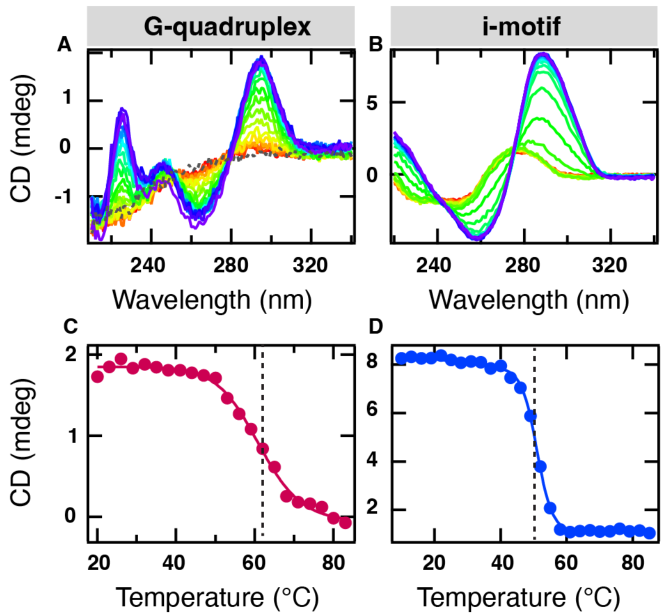


**Figure S13.** Representative far-UV CD spectra of 2 µM Cy5.5-labeled **(A)** G4 in KPhos buffer (10 mM potassium phosphate, 200 mM KCl, 1 mM $K_2EDTA$) at pH 7.0 and **(C)** iM in KPhos buffer pH 6.0. Thermal denaturation of Cy5.5-labeled **(C)** G4 monitored by CD at 295 nm and **(D)** iM monitored by CD at 287 nm. Continuous line overlaid on thermal melts represent a fit to Equation 1. Dashed line represents the melting temperature ($T_m$) extracted from the thermal fits.

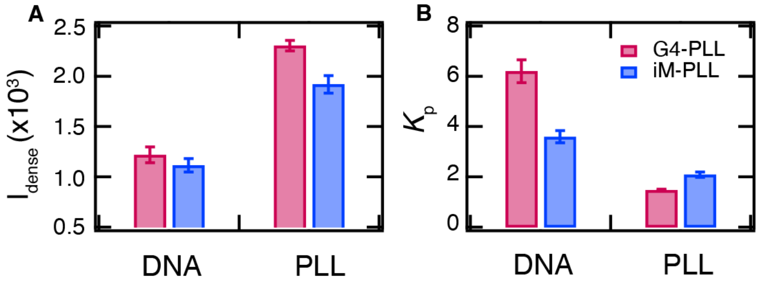


**Figure S14**. Differences between Cy5.5-labeled G4/iM and FITC-PLL partitioning into condensates. **(A)** Average fluorescence intensities of the dense phase for mixtures of G4/PLL and iM/PLL samples in KPhos buffer (10 mM potassium phosphate, 200 mM KCl, 1 mM $K_2$EDTA pH 7.0). **(B)** Partitioning coefficient ($K_p$) of DNA and PLL calculated from fluorescence microscopy images of samples containing 5 µM Cy5.5-labeled G4 (pink) or iM (blue) with 0.1 mg/ml FITC-labeled PLL in KPhos buffer. Standard error of the mean calculated of ten condensates and ten dilute regions. Error bars for partitioning coefficient ($K_p$) are derived from propagating the standard error of ten condensates for each respective condition.

**Table S1.** Thermodynamics and Kinetics of FRET-labeled G4 measured in the nucleus of U-2 OS cells. Error is one standard deviation of the fit. Average error is the standard error.

| Cell | $T_M$ (°C) | ΔH (kcal/mol) | $\tau_1$ (s) | $\tau_2$ (s) | β |
|---|---|---|---|---|---|
| 1 | 44.3 ± 0.7 | 19.7 ± 0.7 | | 11 ± 1 | 0.81 ± 0.02 |
| 2 | 54.0 ± 0.5 | 25 ± 1 | | 1.92 ± 0.05 | 0.83 ± 0.03 |
| 3 | 48.5 ± 0.4 | 32 ± 2 | | | |
| 4 | 43.3 ± 0.5 | 32 ± 3 | | | |
| 5 | 43 ± 1 | 36 ± 3 | | | |
| 6 | 48.8 ± 0.8 | 15 ± 1 | | | |
| 7 | 47.0 ± 0.8 | 25 ± 2 | 0.23 ± 0.03 | 3.0 ± 0.1 | 1.43 ± 0.09 |
| 8 | 44.2 ± 0.9 | 24 ± 1 | 0.48 ± 0.05 | 6.2 ± 0.3 | 1 ± 0 |
| 9 | 42.7 ± 0.4 | 27.3 ± 0.7 | | 14.5 ± 0.6 | 1.41 ± 0.04 |
| 10 | 49.8 ± 0.2 | 40 ± 2 | | | |
| 11 | 47.2 ± 0.3 | 29.2 ± 0.9 | | 0.91 ± 0.07 | 0.55 ± 0.03 |
| 12 | 48.0 ± 0.5 | 27 ± 1 | | | |
| 13 | 36.5 ± 0.1 | 42 ± 1 | | 3.6 ± 0.1 | 0.74 ± 0.03 |
| 14 | 34.6 ± 0.3 | 58 ± 5 | | 8.5 ± 0.6 | 0.72 ± 0.02 |
| 15 | 33.1 ± 0.2 | 49 ± 2 | | 7.4 ± 0.6 | 0.79 ± 0.04 |
| 16 | 39.3 ± 0.6 | 41 ± 4 | | | |
| 17 | 41.1 ± 0.7 | 42 ± 4 | | | |
| 18 | 47.9 ± 0.6 | 28 ± 1 | 0.34 ± 0.06 | 10.6 ± 0.3 | 1 ± 0 |
| 19 | 45.5 ± 0.6 | 32 ± 2 | 0.18 ± 0.09 | 11.7 ± 0.7 | 1 ± 0 |
| 20 | 44.6 ± 0.8 | 23 ± 1 | 0.38 ± 0.08 | 11.6 ± 0.9 | 1.2 ± 0.1 |
| 21 | 44.5 ± 0.4 | 32 ± 2 | 0.41 ± 0.09 | 14 ± 3 | 1.2 ± 0.2 |
| 22 | 45.1 ± 0.8 | 57 ± 8 | | | |
| 23 | 43.9 ± 0.8 | 45 ± 4 | | | |
| 24 | 55.7 ± 0.5 | 25 ± 1 | | | |
| 25 | 38.0 ± 0.9 | 43 ± 5 | | | |
| 26 | 54.5 ± 0.6 | 27 ± 2 | 1.0 ± 0.1 | 4.9 ± 0.4 | 2.3 ± 0.6 |
| 27 | 49.2 ± 0.5 | 25 ± 1 | | | |
| 28 | 38 ± 1 | 38 ± 4 | | | |
| 29 | 42.8 ± 0.6 | 37 ± 2 | | 2.2 ± 0.1 | 0.79 ± 0.04 |
| 30 | 50.1 ± 0.4 | 24.7 ± 0.9 | | 4.9 ± 0.5 | 1 ± 0 |
| 31 | 50.0 ± 0.5 | 28 ± 2 | | | |
| 32 | 45.8 ± 0.3 | 37 ± 2 | | | |
| 33 | 39.5 ± 0.6 | 70 ± 1 | | | |
| 34 | 45.5 ± 0.4 | 35 ± 2 | | | |
| 35 | 42.8 ± 0.7 | 73 ± 1 | | | |
| 36 | 46.5 ± 0.4 | 35 ± 2 | | | |
| **AVERAGE** | **44.9 ± 0.9** | **36 ± 2** | **0.43 ± 0.07** | **7 ± 1** | **1.0 ± 0.1** |

**Table S2.** Thermodynamics of FRET-labeled G4 measured in vitro. Error is one standard deviation of the fit. Average error is the standard error.

| | **Trial 1** | | **Trial 2** | | **Trial 3** | | **Average** | |
|---|---|---|---|---|---|---|---|---|
| **Buffer** | **$T_M$ (°C)** | **ΔH (kcal/mol)** | **$T_M$ (°C)** | **ΔH (kcal/mol)** | **$T_M$ (°C)** | **ΔH (kcal/mol)** | **$T_M$ (°C)** | **ΔH (kcal/mol)** |
| KPhos pH 7.0 | 54.8 ± 0.2 | 26.5 ± 0.6 | 54.2 ± 0.2 | 25.8 ± 0.6 | 54.9 ± 0.4 | 24 ± 1 | **54.6 ± 0.2** | **25.4 ± 0.7** |
| DPBS pH 7.0 | 27.4 ± 0.3 | 21.0 ± 0.5 | 28.4 ± 0.5 | 19.0 ± 0.7 | 34 ± 1 | 15.3 ± 0.9 | **30 ± 2** | **18 ± 2** |
| 100 mg/ml PEG | 56.1 ± 0.2 | 29.1 ± 0.5 | 55.7 ± 0.2 | 29.7 ± 0.7 | 56.2 ± 0.2 | 28.6 ± 0.8 | **56.0 ± 0.2** | **29.1 ± 0.3** |
| 200 mg/ml PEG | 58.8 ± 0.2 | 32.2 ± 0.9 | 58.6 ± 0.3 | 32 ± 1 | 59.3 ± 0.2 | 32.3 ± 0.8 | **58.9 ± 0.2** | **32.2 ± 0.1** |
| 300 mg/ml PEG | 63.4 ± 0.1 | 42.7 ± 0.4 | 63.4 ± 0.1 | 42.7 ± 0.4 | 64.0 ± 0.2 | 38.9 ± 0.9 | **63.6 ± 0.2** | **41 ± 1** |
| 25% NER | 49.5 ± 0.2 | 30.2 ± 0.5 | 47.9 ± 0.1 | 29.2 ± 0.3 | 48.3 ± 0.2 | 27.8 ± 0.4 | **48.6 ± 0.5** | **29.1 ± 0.7** |
| 50% NER | 45.6 ± 0.4 | 27.8 ± 0.8 | 45.8 ± 0.3 | 27.4 ± 0.5 | 45.5 ± 0.2 | 28.7 ± 0.5 | **45.6 ± 0.1** | **28.0 ± 0.4** |
| 75% NER | 42.5 ± 0.5 | 26 ± 1 | 42.3 ± 0.4 | 25.6 ± 0.7 | 41.8 ± 0.2 | 27.2 ± 0.7 | **42.2 ± 0.2** | **26.3 ± 0.5** |
| 100% NER | 36 ± 1 | 13.3 ± 0.7 | 43 ± 2 | 11.0 ± 0.5 | 36 ± 1 | 12.8 ± 0.5 | **38 ± 2** | **12.4 ± 0.7** |
| 10 mg/ml ssDNA | 50.7 ± 0.5 | 31 ± 2 | 50.6 ± 0.5 | 32 ± 2 | 48 ± 1 | 33 ± 3 | **49.8 ± 0.9** | **32.0 ± 0.6** |
| 1 mg/ml BSA* | 49.2 ± 0.3 | 34 ± 1 | 49.2 ± 0.2 | 33 ± 0.6 | 49.5 ± 0.1 | 34 ± 0.4 | **49.3 ± 0.1** | **33.7 ± 0.3** |
| Complementary DNA | DNF | DNF | DNF | DNF | DNF | DNF | **DNF** | **DNF** |
| Master Mix | 52.7 ± 0.4 | 38 ± 1 | 51.2 ± 0.6 | 37 ± 2 | 51.6 ± 0.4 | 38 ± 1 | **51.8 ± 0.4** | **37.7 ± 0.3** |
| Cytoplasm Lysate | 48.2 ± 0.5 | 31 ± 1 | 50.9 ± 0.7 | 29 ± 2 | 45.6 ± 0.4 | 31 ± 1 | **48 ± 2** | **30.3 ± 0.7** |
| Nuclear Lysate | 48.6 ± 0.5 | 32 ± 1 | 47.0 ± 0.3 | 33 ± 1 | 46.9 ± 0.6 | 37 ± 3 | **47.5 ± 0.6** | **34 ± 2** |
| Condensates* | DNF | DNF | DNF | DNF | DNF | DNF | **22.5 ± 0.5** | **56 ± 11** |

*Temperature jump data with FReI

**Table S3.** Thermodynamics and Kinetics of FRET-labeled iM measured in the nucleus of U-2 OS cells. Error is one standard deviation of the fit. Average error is the standard error.

| Cell | $T_M$ (°C) | $\Delta H$ (kcal/mol) | $\tau_1$ (s) | $\tau_2$ (s) | $\beta$ |
|---|---|---|---|---|---|
| 1 | 41 ± 2 | 21 ± 4 | | | |
| 2 | 44 ± 1 | 23 ± 4 | 0.15 ± 0.07 | 1.3 ± 0.3 | 1.0 ± 0 |
| 3 | 42.2 ± 0.4 | 35 ± 2 | 0.16 ± 0.03 | 5.4 ± 0.4 | 1.7 ± 0.3 |
| 4 | 42.2 ± 0.5 | 42 ± 4 | | | |
| 5 | 52.9 ± 0.7 | 56 ± 7 | | | |
| 6 | 46.2 ± 0.7 | 21 ± 2 | | | |
| 7 | 45 ± 2 | 21 ± 4 | | | |
| 8 | 51 ± 1 | 20 ± 3 | 0.25 ± 0.02 | 4.5 ± 0.2 | 2.3 ± 0.4 |
| 9 | 43.7 ± 0.5 | 32 ± 3 | | | |
| 10 | 47.7 ± 0.3 | 41 ± 3 | | | |
| 11 | 49 ± 2 | 16 ± 3 | | | |
| 12 | 42.1 ± 0.6 | 26 ± 3 | | | |
| 13 | 40.9 ± 0.8 | 46 ± 7 | 0.24 ± 0.03 | 5.3 ± 0.2 | 2.4 ± 0.3 |
| 14 | 42 ± 1 | 29 ± 5 | | | |
| 15 | 51.0 ± 0.6 | 36 ± 3 | 0.30 ± 0.03 | 5.1 ± 0.2 | 3.0 ± 0.5 |
| 16 | 53.3 ± 0.6 | 39 ± 3 | | | |
| 17 | 44.0 ± 0.7 | 43 ± 5 | | | |
| 18 | 53 ± 1 | 37 ± 8 | | | |
| 19 | 46 ± 2 | 32 ± 5 | | | |
| 20 | 46.5 ± 0.5 | 42 ± 4 | | | |
| 21 | 44.0 ± 0.4 | 27 ± 1 | | | |
| 22 | 44 ± 1 | 33 ± 7 | | | |
| 23 | 47 ± 1 | 22 ± 3 | | | |
| 24 | 45.5 ± 0.5 | 28 ± 2 | | | |
| 25 | 52 ± 2 | 22 ± 3 | | | |
| 26 | 48 ± 3 | 24 ± 4 | | | |
| 27 | 50 ± 2 | 28 ± 6 | | | |
| 28 | 43.5 ± 0.7 | 31 ± 3 | | | |
| 29 | 50 ± 2 | 30 ± 7 | | | |
| 30 | 49.8 ± 0.9 | 30 ± 7 | 0.28 ± 0.03 | 5.4 ± 0.7 | 1 ± 0 |
| 31 | 59 ± 2 | 28 ± 5 | | | |
| 32 | 43.9 ± 0.9 | 43 ± 7 | | | |
| 33 | 43 ± 1 | 28 ± 4 | | | |
| 34 | 47.8 ± 0.9 | 33 ± 4 | | | |
| 35 | 51 ± 2 | 31 ± 6 | | | |
| 36 | 41.9 ± 0.4 | 72 ± 8 | 0.21 ± 0.03 | 4.1 ± 0.4 | 1.5 ± 0.4 |
| 37 | 42 ± 1 | 30 ± 4 | 0.25 ± 0.02 | 5.5 ± 0.7 | 4 ± 2 |
| 38 | 49.3 ± 0.8 | 130 ± 50 | | | |
| 39 | 39 ± 1 | 72 ± 2 | | | |
| 40 | 46.4 ± 0.4 | 50 ± 4 | 0.19 ± 0.03 | 3.6 ± 0.2 | 2.4 ± 0.4 |
| 41 | 52.7 ± 0.8 | 30 ± 3 | 0.25 ± 0.02 | 4.0 ± 0.1 | 7 ± 2 |
| 42 | 41.2 ± 0.8 | 24 ± 3 | | | |

| | | | | | |
|---|---|---|---|---|---|
| 43 | 41 ± 1 | 23 ± 2 | 0.18 ± 0.02 | 3.9 ± 0.5 | 2 ± 1 |
| 44 | 50.8 ± 0.5 | 56 ± 7 | | | |
| 45 | 47.3 ± 0.7 | 25 ± 3 | | | |
| 46 | 46.0 ± 0.6 | 22 ± 2 | | | |
| 47 | 55 ± 2 | 20 ± 2 | | | |
| 48 | 46 ± 1 | 22 ± 4 | | | |
| 49 | 46 ± 1 | 31 ± 5 | | | |
| 50 | 51 ± 2 | 24 ± 4 | | | |
| 51 | 45.9 ± 0.8 | 36 ± 4 | | | |
| 52 | 51 ± 2 | 24 ± 4 | 0.27 ± 0.02 | 11.5 ± 0.3 | 6 ± 1 |
| 53 | 44 ± 1 | 32 ± 4 | | | |
| 54 | 47.9 ± 0.6 | 42 ± 5 | | | |
| 55 | 51 ± 4 | 20 ± 7 | | | |
| 56 | 36.0 ± 0.5 | 41 ± 5 | | | |
| 57 | 44 ± 1 | 25 ± 2 | | | |
| 58 | 51 ± 3 | 30 ± 9 | | | |
| 59 | 41 ± 1 | 36 ± 8 | | 8.7 ± 0.4 | 5 ± 1 |
| 60 | 40.8 ± 0.6 | 43 ± 5 | 0.24 ± 0.03 | 8.7 ± 0.3 | 2.4 ± 0.3 |
| **AVERAGE** | **46.5 ± 0.6** | **34 ± 2** | **0.23 ± 0.01** | **5.5 ± 0.7** | **3.0 ± 0.5** |

**Table S4.** Thermodynamics of FRET-labeled iM measured in vitro. Error is one standard deviation of the fit. Average error is the standard error. DNF where data could not be fit.

| | **Trial 1** | | **Trial 2** | | **Trial 3** | | **Average** | |
|---|---|---|---|---|---|---|---|---|
| **Buffer** | **$T_M$ (°C)** | **ΔH (kcal/mol)** | **$T_M$ (°C)** | **ΔH (kcal/mol)** | **$T_M$ (°C)** | **ΔH (kcal/mol)** | **$T_M$ (°C)** | **ΔH (kcal/mol)** |
| KPhos pH 7.0 | 27.4 ± 0.8 | 110 ± 40 | 27.1 ± 0.7 | 150 ± 70 | 28.1 ± 0.3 | 90 ± 10 | **27.5 ± 0.3** | **120 ± 20** |
| DPBS pH 7.0 | 30.8 ± 0.5 | 73 ± 10 | 30 ± 1 | 30 ± 6 | 31 ± 2 | 15 ± 3 | **30 ± 2** | **39 ± 20** |
| 100 mg/ml PEG | 27.3 ± 0.1 | 77 ± 3 | 29.3 ± 0.2 | 100 ± 7 | 29.1 ± 0.3 | 100 ± 10 | **28.5 ± 0.7** | **92 ± 8** |
| 200 mg/ml PEG | 27.1 ± 0.2 | 87 ± 6 | 29.3 ± 0.2 | 96 ± 7 | 29.3 ± 0.3 | 100 ± 10 | **28.6 ± 0.7** | **94 ± 4** |
| 300 mg/ml PEG | 28.4 ± 0.3 | 110 ± 20 | 31.1 ± 0.1 | 86 ± 4 | 29.5 ± 0.3 | 77 ± 9 | **29.7 ± 0.8** | **90 ± 10** |
| 25% NER | 31 ± 2 | 30 ± 8 | 25.8 ± 0.4 | 60 ± 6 | DNF | DNF | **28 ± 2** | **45 ± 20** |
| 50% NER | 24.2 ± 0.3 | 76 ± 9 | DNF | DNF | DNF | DNF | **24.2 ± 0.3** | **76 ± 9** |
| 75% NER | DNF | DNF | DNF | DNF | DNF | DNF | **DNF** | **DNF** |
| 100% NER | DNF | DNF | DNF | DNF | DNF | DNF | **DNF** | **DNF** |
| 10 mg/ml ssDNA | 29.8 ± 0.2 | 100 ± 10 | 30.3 ± 0.2 | 94 ± 8 | 32.3 ± 0.6 | 86 ± 20 | **30.8 ± 0.8** | **93 ± 4** |
| 1 mg/ml BSA | 27.0 ± 0.3 | 100 ± 15 | 26.6 ± 0.3 | 100 ± 14 | 30.0 ± 0.2 | 95 ± 6 | **28 ± 2** | **98 ± 3** |
| Complementary DNA | 27.2 ± 0.2 | 86 ± 6 | 28.4 ± 0.2 | 86 ± 6 | 28.3 ± 0.2 | 76 ± 6 | **28.0 ± 0.4** | **83 ± 4** |
| Master Mix | 31.8 ± 0.3 | 91 ± 10 | 31.2 ± 0.2 | 66 ± 3 | 29.7 ± 0.4 | 68 ± 7 | **30.9 ± 0.6** | **75 ± 8** |
| Cytoplasm Lysate* | 38.0 ± 0.6 | 16 ± 1 | 35 ± 1 | 17 ± 2 | DNF | DNF | **37 ± 1** | **16.5 ± 0.5** |
| Nuclear Lysate** | 47.4 ± 0.4 | 19.0 ± 0.7 | 49.4 ± 0.6 | 18 ± 1 | 39.3 ± 0.4 | 18 ± 0.8 | **44 ± 3** | **17.5 ± 0.9** |
| Condensates*** | 30.4 ± 0.3 | 74 ± 10 | DNF | DNF | DNF | DNF | **30.4 ± 0.3** | **74 ± 10** |

*Only two measurements due to limited iM sample

**Fourth trial: Tm = 37.8 ± 0.9 °C and ΔH = 15 ± 1 kcal/mol

***Temperature jump data collected with FReI

**Table S5.** FRET-labeled G4 kinetics measured in vitro. Relaxation lifetimes ($\tau_2$) and $\beta$ factor following a temperature jump to $T_M$. Error is one standard deviation of the fit. Average error is the standard error.

| | **Trial 1** | | **Trial 2** | | **Trial 3** | | **Average** | |
|---|---|---|---|---|---|---|---|---|
| **Buffer** | $\tau_2$ **(s)** | $\beta$ | $\tau_2$ **(s)** | $\beta$ | $\tau_2$ **(s)** | $\beta$ | $\tau_2$ **(s)** | $\beta$ |
| KPhos pH 7.0* | 0.24 ± 0.02 | 1 | 0.23 ± 0.02 | 1 | 0.13 ± 0.01 | 1 | **0.19 ± 0.03** | **1** |
| 200 mg/ml PEG | 0.87 ± 0.09 | 2.2 ± 0.7 | | | | | | |
| 50% NER** | 0.30 ± 0.01 | 1.9 ± 0.1 | 0.198 ± 0.008 | 2.2 ± 0.2 | 0.15 ± 0.03 | 1 | **0.23 ± 0.04** | **1.7 ± 0.3** |
| 10 mg/ml ssDNA*** | 0.27 ± 0.06 | 2.1 ± 0.9 | 0.23 ± 0.05 | 1.4 ± 0.3 | 0.33 ± 0.03 | 1 | **0.27 ± 0.02** | **1.6 ± 0.2** |
| Nuclear Lysate**** | 0.20 ± 0.02 | 1.8 ± 0.3 | 0.27 ± 0.03 | 1 | 0.226 ± 0.008 | 1 | **0.23 ± 0.01** | **1.5 ± 0.3** |

*Fourth trial: $\tau_2$ 0.14 ± 0.02 and $\beta$ 1; with $\beta$ resulted in overfitting (high errors), so $\beta$ held to one for all fits
**Fourth trial: $\tau_2$ 0.28 ± 0.01 and $\beta$ 1.50 ± 0.07; where $\beta$ resulted in overfitting (high errors), $\beta$ held to one
***Fourth trial: $\tau_2$ 0.24 ± 0.04 and $\beta$ 1.9 ± 0.4; where $\beta$ resulted in overfitting (high errors), $\beta$ held to one
****Fourth trial: $\tau_2$ 0.23 ± 0.03 and $\beta$ 2.3 ± 0.7; where $\beta$ resulted in overfitting (high errors), $\beta$ held to one

**Table S6.** FRET-labeled iM kinetics measured in vitro. Relaxation lifetimes ($\tau_2$) and $\beta$ factor following a temperature jump to $T_M$. Error is one standard deviation of the fit. Average error is the standard error.

| | **Trial 1** | | **Trial 2** | | **Trial 3** | | **Average** | |
|---|---|---|---|---|---|---|---|---|
| **Buffer** | **$\tau_2$ (s)** | **$\beta$** | **$\tau_2$ (s)** | **$\beta$** | **$\tau_2$ (s)** | **$\beta$** | **$\tau_2$ (s)** | **$\beta$** |
| KPhos pH 7.0* | 26.3 ± 0.7 | 2.1 ± 0.2 | 19.8 ± 0.4 | 1.75 ± 0.08 | 29.1 ± 0.6 | 1.02 ± 0.04 | **24 ± 2** | **1.5 ± 0.3** |
| 200 mg/ml PEG | 34.6 ± 0.1 | 1.141 ± 0.007 | 26.1 ± 0.1 | 1.045 ± 0.008 | 28.5 ± 0.1 | 1.106 ± 0.009 | **30 ± 3** | **1.10 ± 0.02** |
| 10 mg/ml ssDNA** | 14.76 ± 0.05 | 0.098 ± 0.005 | 14.37 ± 0.07 | 1.010 ± 0.007 | 26.3 ± 0.3 | 0.95 ± 0.01 | **28 ± 7** | **0.95 ± 0.03** |
| Nuclear Lysate | 34.2 ± 0.1 | 0.755 ± 0.003 | 35.12 ± 0.09 | 0.784 ± 0.002 | 19.31 ± 0.06 | 0.985 ± 0.005 | **30 ± 5** | **0.84 ± 0.07** |

*Fourth trial: $\tau_2$ 22.5 ± 0.5 and $\beta$ 1.18 ± 0.05

**Fourth trial: $\tau_2$ 43.7 ± 0.1 and $\beta$ 0.875 ± 0.007

**Table S7.** List of unlabeled and labeled G4 and iM sequences.

| Oligonucleotide sequence (5′ to 3′) | |
|---|---|
| unlabeled G4 | GGT TGG TGT GGT TGG |
| G4 | TAMRA-GGT TGG TGT GGT TGG-FAM |
| unlabeled iM | TAA CCC CCT AAC CCC CTA ACC CCC TAA CCC CCT AA |
| iM | TAMRA-TAA CCC CCT AAC CCC CTA ACC CCC TAA CCC CCT AA-FAM |
| G4-Cy5.5 | Cy5.5-GGT TGG TGT GGT TGG |
| iM-Cy5.5 | TAA CCC CCT AAC CCC CTA ACC CCC TAA CCC CCT AA-Cy5.5 |
| G4 complement | CCA ACC ACA CCA ACC |
| iM complement | TTA GGG GGT TAG GGG GTT AGG GGG TTA GGG GGT TA |
